# Macrophages heterogeneity and significance during human fetal pancreatic development

**DOI:** 10.1101/2023.07.27.549142

**Authors:** Adriana Migliorini, Sabrina Ge, Michael H. Atkins, Rangarajan Sambathkumar, Angel Sing, Conan Chua, Adam J. Gehring, Gordon M. Keller, Faiyaz Notta, Maria Cristina Nostro

## Abstract

Organogenesis is a complex process that relies on a dynamic interplay between extrinsic factors originating from the microenvironment and intrinsic factors specific to the tissue. For the endocrine cells of the islet of Langerhans, the local microenvironment consists of various cell types including pancreatic acinar and ductal cells as well as neuronal, immune, endothelial, and stromal cells. Interestingly, hematopoietic cells have been detected in human pancreas as early as 6 post-conception weeks (PCW)^1,2^, but whether they play a role during islet formation in humans remains largely unknown. To shed light on this question, we performed single nuclei RNA sequencing of the human fetal pancreas during the early weeks of the second trimester, specifically focusing on the molecular interaction between the hematopoietic niche and the pancreatic epithelium. Our analysis identified a wide range of hematopoietic cells as well as two distinct subsets of macrophages that are unique to the fetal pancreas and absent in neonatal or adult pancreatic tissues. Leveraging this discovery, we developed a co-culture system of hESC-derived endocrine-macrophage organoids to model their interaction *in vitro*. Remarkably, we found that macrophages promoted the differentiation and viability of developing endocrine cells *in vitro* and enhanced tissue engraftment in immunocompromised mice, supporting a role for these cells in future tissue engineering strategies for diabetes.

## INTRODUCTION

Human pancreatic development is a complex process orchestrated by a network of varying cell types that constitute the pancreatic niche. While the crosstalk between pancreatic epithelium and surrounding cell types, such as mesenchymal and vascular populations, has been extensively studied^3-7^, little attention has been placed on understanding whether the immune cells support pancreatic organogenesis. During fetal development, immune cells migrate and colonize the fetal organs to establish peripheral tolerance, a process that is impaired in many autoimmune diseases, such as type 1 diabetes (T1D)^8^.

Alongside the establishment of immune tolerance, several lines of evidence suggest that immune cells, and specifically macrophages, are critical for organogenesis^9,10^. Precisely, mouse studies have demonstrated that stroma-associated macrophages and islet-associated macrophages might have different origins and properties^11^ and the depletion of CSF1-dependent macrophages (observed in *Csf1^op^/Csf1^op^* mice) affects the morphology and composition of developing islets^12^. Additional studies have further shown that CCR2+ macrophages may be required for islet proliferation and maturation during the perinatal time^13^. However, challenges associated with studying human development have precluded the full understanding of their role during human pancreas development. Here, we provide evidence supporting a model where embryonic macrophages play a broader role in pancreas physiology and organogenesis than previously thought. To this aim, we molecularly characterized the hematopoietic cells in the developing pancreas by performing single nuclei RNA sequencing (snRNAseq) during the second trimester of human gestation. This analysis resulted in the identification of 30 unique cell clusters describing the pancreatic epithelium and its microenvironment, comprising 6 distinct hematopoietic cell types. Interestingly, within the hematopoietic niche, we identified 2 populations of macrophages that were distinguished by the expression of *TIMD4*, *CD36* and *LYVE1*. We uncovered that both types of macrophages shared similarities with regulatory macrophages and differ from the macrophage populations previously identified in the human postnatal and adult pancreas^14^. To explore the potential role of fetal-like macrophages during human pancreatic development, we developed a co-culture system using human embryonic stem cells (hESCs). By modeling yolk sac (YS) hematopoiesis and pancreatic development, we generated hESC-derived embryonic macrophages (eMAC)^15^ and pancreatic endoderm^16^ that closely resembled their fetal counterparts in terms of cellular composition and molecular profiles”. Remarkably, we determined that hESC-derived eMAC improved the development and viability of hESC-derived endocrine cells when compared to standard growth conditions. Finally, we found that transplantation of eMAC-endocrine co-cultures improved graft vascularization, insulin secretion, and the frequency of hormone+ cells in the murine subcutaneous space. This comprehensive interrogation of the hematopoietic cells within the pancreatic niche and its applications could pave the way for novel stem cell-based transplantation modalities and tissue engineering strategies for diabetes.

## RESULTS

### Single-cell transcriptomic analysis of the human fetal pancreas

To map the cellular and molecular states of the developing human pancreas and its niche, we performed single nuclei RNA sequencing (snRNAseq) of fetal pancreas collected between 14 to 18 PCW (Figure S1A), a period characterized by prominent tissue remodeling and islet morphogenesis^17^ (Figure 1A). We isolated and enriched intact nuclei using Fluorescence Activated Cell Sorting (FACS) (Figure S1 B-D). In total, 73,274 (n=6 distinct tissue donors) pancreatic cell nuclei passed quality control and doublet exclusion. Graph-based Leiden clustering determined major cell classes, including epithelial, mesenchymal, endothelial, neuronal, erythroid, and immune populations (Figure 1B). We applied a descriptive nomenclature based on the identified differentially expressed genes (DEGs) to annotate the 30 molecularly distinct cell clusters (Figure 1B,D). While all cell states were found throughout the three developmental periods analyzed (Figure S2A,B), frequencies and cell numbers varied among gestational ages (Figure 1C). Sex-related differences among samples were not detected (Figure S2C). Intriguingly, cell cycle scoring analysis highlighted that the cells found in S or G2M phase were either acinar, ductal, mesenchymal or erythroid in nature, while most of the remaining fetal pancreatic cells were found in G1 phase (Figure S2D,E).

**Figure 1.**
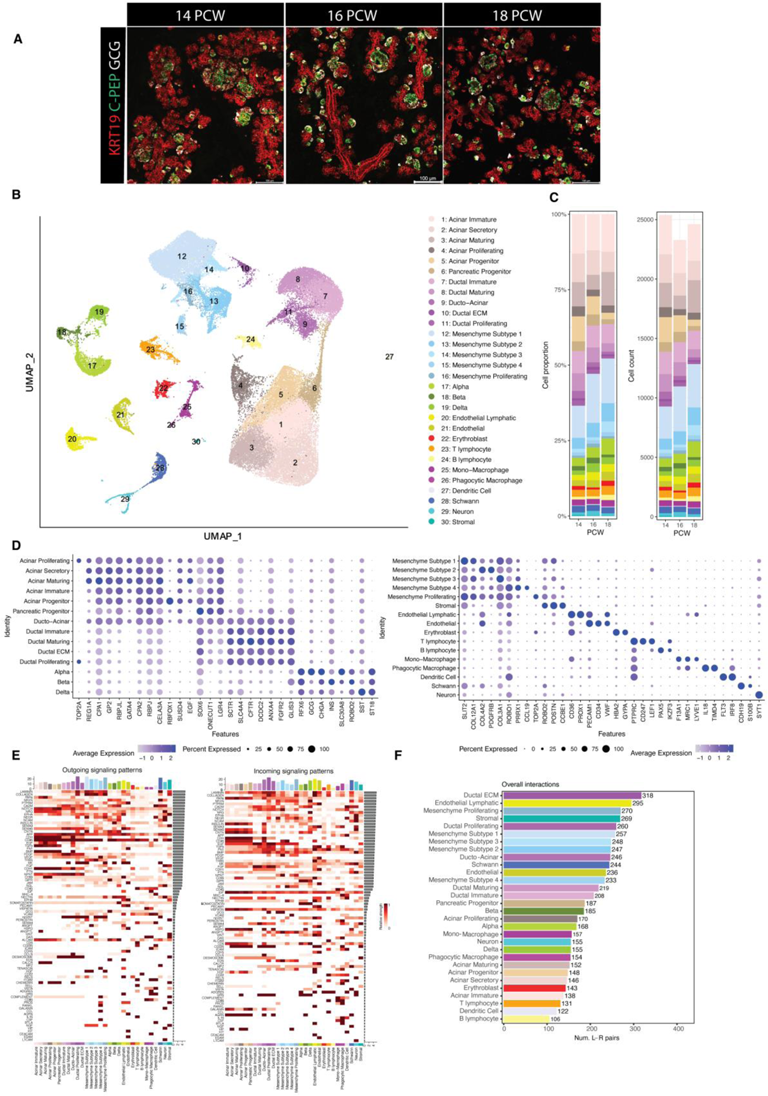
Single cells transcriptomic phenotyping of the human fetal pancreas. **A**, Confocal Laser Scanning Microscopy (CLSM) images of fetal pancreatic epithelium across three Post Conception Weeks (14, 16, 18 PCW) stained for cytokeratin 19 (KRT19+, red), c-peptide (C-PEP, green) and glucagon (GCG, white). **B**, UMAP visualization of 14-18 PCW fetal pancreas samples (n=6, with 2 biologically independent samples per PCW). Cells are clustered by their major cell class, with distinct colors and numbers representing cluster assignments. **C**, Fetal pancreas composition (frequency and cell numbers) by PCW. Colors correspond to the cell clusters featured in **B**. **D**, Dot plot showing gene expression of cell cluster markers. **E**, Heatmap summarizing significant signaling pathway interactions by ligand-receptor analysis (L-R) across all the major cell types of the fetal pancreas. **F**, Bar plot of the overall interactions detected by L-R analysis within all the major cell types in the fetal pancreas.

Glycoprotein 2 (GP2)^+^ fetal acinar cells accounted for the majority of the developing pancreatic epithelium in the early second trimester of gestation^18,19^ (Figure S3A). To resolve acinar cell heterogeneity, we performed Gene Set Enrichment Analysis (GSEA) and identified known and novel gene markers labeling different acinar states, which could be further subsetted into 4 developmentally distinct acinar clusters: Acinar Immature (*ROR1*^high^/*REG1A*^low^)^20,21^, Acinar Secretory (*UNC13B*^high^/*TM4SF4*^high^)^22,23^, Acinar Maturing (*EGF*^high^/*CPA1*^high^)^24,25^ and Acinar proliferating (*TOPA2*^high^) (Figure S3B-D). Additionally, using ligand-receptor (L-R) pairing analysis, we explored acinar-derived niche cues. Consistent with previous studies, our analysis suggested that EGF^26,27^, TGFβ^28^ and BMP^29^ signalling networks participate in the pancreatic epithelium cross-talk during the second trimester of human development^16,30^ (Figure S3E). Interestingly, our analysis revealed that the acinar cells are the major source of ligands specific for TGFβ signaling to macrophages, endothelial and T cells (Figure S3E middle panel).

To map ductal cell diversity, we performed GSEA analysis and characterized the different ductal cell states (Figure S4A). We identified 4 molecularly distinct ductal clusters: Ductal Immature (*COL27A1*^high^/*CFTR*^low^)^31,32^, Ductal Maturing (*CFTR*^high^/*PPARGC1A*^high^)^31,33^, Ducto-Acinar (*CAMK1D*^high^/*GP2*^+^)^33,34^, and Ductal ECM (*ZEB1*^+^/*SLIT2*^high^)^35,36^ (Figure S4B,C). The ducto-acinar cluster, while predominantly ductal in nature, also express the acinar genes *CPA1* and *GP2* (Figure 1D and Figure S4D). The existence of ductal cells expressing acinar markers was validated in sections of fetal pancreas by immunostaining for GP2 (acinar marker) and KRT19 (ductal marker) (Figure S4E). These data provide a higher-order map and a catalogue of molecular profiles of exocrine cell types within the developing human fetal pancreas epithelium.

As previously described in the developing mouse and human pancreas^37,38^, we detected a pronounced variety of mesenchymal cells (Figure S5A,B). We catalogued a stromal cluster^39^ (Figure 1B,D) as well as 5 molecularly distinct mesenchyme clusters that include the known *PBX1*^high^/*POSTN* ^high^ mesenchyme subpopulation^4^ (mesenchyme subtype 1) as well as subpopulations characterized as *MYO1B*^high^/*INPP4B*^high^ (mesenchyme subtype 2), *MEOX2*^high^/*EPHB2*^high^ (mesenchyme subtype 3), *CCL21*^+^/*IL34*^+^ (mesenchyme subtype 4) and *TOP2A*^+^/*EZH2*^+^ (mesenchyme proliferating) (Figure S5C). Based on L-R pairing analysis, the mesenchyme clusters, neuronal and stromal clusters represent the major source of the FGF signalling network in the developing pancreas (Figure S5D). Interestingly, we found that only mesenchyme subtype 4, transcriptionally resembling a fibroblastic reticular cell phenotype^40^, specifically interacts with fetal pancreatic T cells and macrophages via CCL21 and IL34^41^, respectively (Figure S5E,F). IL34 and CSF1 are the two colony-stimulating factor 1 receptor (CSF1R) ligands, which are essential to the development and maintenance of microglia in the embryonic and adult brain^42,43^.

We next sought to phenotype the fetal endocrine pancreatic cells by performing sub-clustering of the endocrine cells detected in Figure 1B (Figure S6A,B). We identified three subpopulations of alpha cells (early, middle, and late) and two subpopulations of delta cells (early and late), likely reflecting their differentiation status, as their frequencies and cell numbers varied among gestational age (Figure S6C). Further, we conducted pseudo-time analysis and integrated DEG to annotate cellular states and differentiation trajectories of the developing pancreatic endocrine cells (Figure S6D,E). Intriguingly, pseudo-time analysis validated the notion that alpha cells develop earlier than beta cells in humans^44^ and revealed the dynamic expression of biologically interesting genes such as *DPP4* and *GDA* in alpha cells^45^, *LEPR* and *ERBB4* in delta cells^46^, and *SLCA30A8* and *IAPP* in beta cells in the second trimester (Figure S6F).

To investigate the cell-to-cell communication network events among all the major cell types in the fetal pancreas, we performed L-R pairing analysis and identified 436 significant ligand-receptor pairings, representative of 88 signaling pathways (Figure 1E). Confirming the relevance of this dataset, we identified signaling pathways that have previously been recognized to play a significant role during pancreas development (such as Laminin, Collagen, Notch, WNT, FGF, EGF, BMP and GIPR)^47-49^ ^50^ as well as signalling pathways that have been associated with poor prognosis in pancreatic cancer and may have novel relevance in pancreas development, such as cell adhesion molecules part of the NECTIN family^51^ and progranulin encoded by *GRN^52^*. When looking at the overall number of L-R pairing interactions, immune cells accounted for a large proportion of the L-R pairing interactions, underlining an active role of these cells within the pancreatic developmental niche (Figure 1F).

This fetal pancreas dataset can be explored using an interactive web portal (link provided upon acceptance of the manuscript).

### Fetal hematopoietic pancreatic niche

To characterize the hematopoietic cells within the fetal pancreatic niche at 14, 16 and 18 PCW, we performed sub-clustering of the hematopoietic cells identified in our initial analysis (Figure 1B, Figure 2A insert). The frequency of these cells was then calculated for each developmental time analyzed (Figure 2B). We discovered 13 hematopoietic clusters and manually annotated these hematopoietic cell types based on their DEGs and specific gene expression markers (Figure 2A-C).

**Figure 2.**
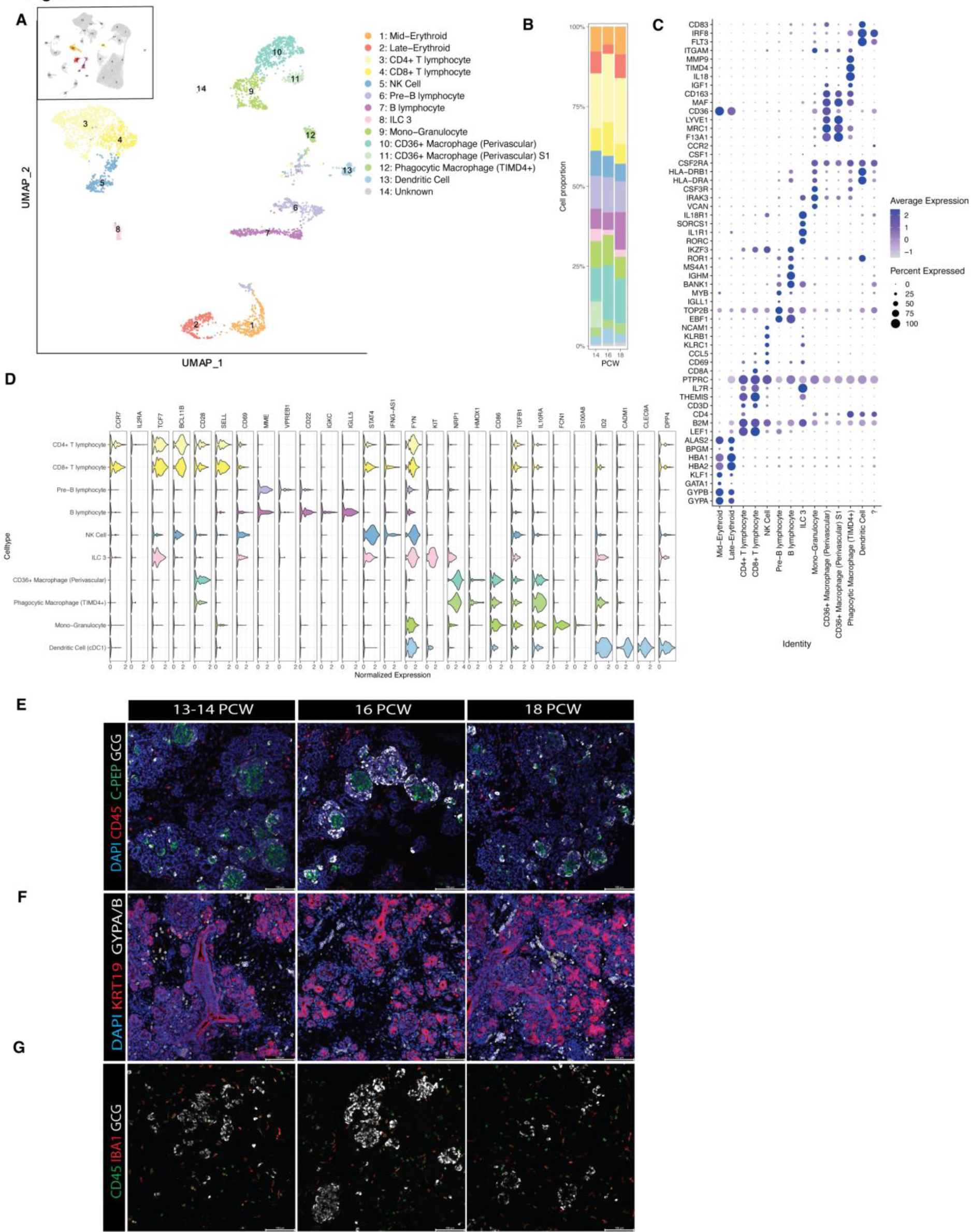
Characterization of the fetal pancreatic hematopoietic niche. **A**, UMAP visualization and cell annotation of fetal pancreatic hematopoietic cells from Figure 1B. Inset UMAP plot in the top left corner highlights the cell subset used while unselected cells are gray. **B**, Pancreatic hematopoietic cell composition (frequency) by PCW. Colors indicate cell clusters as shown in **A**. **C**, Dot plot showing hematopoietic cell marker gene expression. **D**, Violin plots depict the expression distribution of discrete genes in pancreatic hematopoietic cells. **E**, CLSM images of fetal tissue section at 13-18 PCW highlighting CD45+ hematopoietic cells (red) and their proximity to endocrine c-peptide+ (green) and glucagon+ (white) cells. **F**, CLSM images of fetal tissue section at 13-18 PCW highlighting KRT19+ (red) pancreatic cells and GYPA/B+ (white) erythroid cells. **G**, CLSM images of fetal tissue section at 13-18 PCW highlighting CD45+ immune cells (green), IBA1+ macrophages (red) and endocrine glucagon+ (GCG)cells (white) in the developing pancreas.

Erythroid cells clustered mainly in two groups, Late-Erythroid (*HBA1-2*^high^/*GATA1*^-^/*KLF1*^low^/*BPGM*^+^) and proliferating Mid-Erythroid (*KLF1*^+^/*GATA1*^+^/*BPGM*^low^)^53^ (Figure 2A,C; Figure S7A), suggesting that erythroid maturation occurs in the pancreas and that the fetal pancreas contains a pool of proliferating erythroid progenitors that can contribute to supplementing the erythroid output of the fetal liver.

Lymphoid cells clustered into several groups including natural killer (NK), type 3 innate lymphoid cells (ILC3), B lymphocytes, and CD4+ and CD8+ T lymphocytes (Figure 2A,C). Fetal pancreatic NK cells appear to have an immature phenotype as they express archetypal cell-specific markers such as *NCAM1* (*CD56*)*^54^, CCL5* and *CD69*, as well as important inhibitory receptor-like *KLRC1* and *KLRB1,* characteristic of immature NK cells^55^. Concordantly, they expressed low levels of *SELL* and *CD57,* but high levels of *STAT4* and the long noncoding RNA (lncRNA) *IFNG-AS1^56^*, which suggests that they might be poised to secrete inflammatory cytokines (Figure 2C,D). In addition to NK cells, we identified a cluster of *NCR*^-^ ILC3 cells^57^, innate lymphoid cells characterized by the expression of TCF1 (encoded by *TCF7*), *RORC* and *KIT*. ILC3 cells have been reported to play an important role in the maintenance of gut homeostasis^58,59^, and may display a similar role in the pancreas (Figure 2A,C,D).

Unlike rodents, which depend on the thymic turnover of naïve T cells throughout their lifetime, most human T cell development occurs during the fetal period, emphasizing the importance to study the functional competence of prenatal tissue-resident T cells^60^. In the fetus, T cells have been detected in peripheral tissue as early as 10-11 PCW^61^ and, consistent with what is observed in mucosal tissue, CD4+ and CD8+ T cell clusters were identified by snRNAseq analysis at all time points analyzed (Figure 2A,B). Flow cytometric analysis of human fetal pancreas at 16 PCW demonstrated the presence of naïve (CD3+CD45RO+) and memory (CD3+CD45RA+) T cells (Figure S7B). This is consistent with their expression profile, as both *CD4+* and *CD8+* pancreatic fetal T lymphocytes express high levels of *CCR7, TCF1/7, CD28* and *SELL* (*CD62L*) but low levels of *IL2RA* (*CD25*) and *CD69*, which are characteristic markers of memory T cells and naïve T cells, respectively*^62^* (Figure 2D). The presence of T memory cells demonstrates the potential of fetal pancreatic T cells to respond to antigens, emphasizing that fetal pancreatic development may be a relevant stage to understand the pathogenesis of T1D in pediatric patients.

Among the B cell lineage, we detected cells with two differentiation states: proliferating precursor B (pre-B) cells (Figure 2A, Figure S7A) expressing *IGLL1* and *MYB* transcripts, with low levels of *MS4A1* and *IKFZ3,* and B cells expressing *IGMH* and *BANK1* (Figure 2C,D)^53,63^. The presence of B cells during pancreas development is of importance and suggests that pancreas autoantibody detected in children at risk of developing T1D might be developing already during fetal development.

Myeloid cells clustered mainly in two groups, which were assigned to dendritic cells (DC) and monocyte/macrophages (Figure 2A). Based on the expression of *IRF8, CADM1, CLEC9E* and *DPP4* transcripts, the dendritic cell cluster was cataloged as classical DC (cDC1)^64^ (Figure 2C,D).

Amongst the macrophage cluster, we identified one population of monocyte/granulocyte and two subpopulations of tissue-resident macrophages, which we labelled perivascular macrophages (expressing *LYVE1* and the scavenger receptor *CD36^65^*) and phagocytic macrophages (expressing *MMP9^66^, IL18^67^* and the phagocytic receptor *TIMD4^68^*) (Figure 2A,C). Our analysis also identified a third cluster of macrophages arising from one specimen showing many similarities with the perivascular macrophages cluster. Cells within the monocyte/granulocyte cluster express monocyte-related genes such as *CSF3R*^69^, *FCN1*^70^, and *S100A8*^71^, but lack expression of *CD16* (Figure 2C,D).

Both perivascular (*CD36+, LYVE+*) and phagocytic (*TIMD4+*) macrophages display a gene expression profile similar to adult macrophages, including the presence of *ITGAM* and *CD163* (Figure 2C), but lack *CCR2* expression, a chemokine receptor often found in circulating inflammatory monocytes originating from the definitive hematopoietic program^72,73^. Interestingly, perivascular macrophages express higher levels of *MRC1, CD163,* and *CD28* than phagocytic macrophages (Figure 2C), and when compared to the other immune cells, perivascular macrophages express higher levels of *SPP1, PDGFB, ITGB5, EDNRB*, and *IGFBP4* (data not shown), all genes associated with angiogenesis^74^. We also detected notable expression levels of *NRP1^75^*, *HOMX1^76^,* and *IL-10RA^77^* as well as *CD86* and *TGFB1* within the monocyte/granulocyte and macrophages populations, indicating their active role as antigen-presenting cells and their possible link to a tolerogenic phenotype (Figure 2D). To corroborate and explore the distribution of pancreatic hematopoietic cells detected by snRNAseq, we performed immunostaining and Confocal Laser Scanning Microscopy (CLSM) analysis of fetal pancreatic sections at 14, 16, and 18 PCW. Our analysis detected CD45+ hematopoietic cells (Figure 2E), anucleated GYPA/B+ erythrocytes, as well as nucleated GYPA/B+ erythroid cells (Figure 2F), and IBA1+ macrophages (Figure 2G) at all the developmental stages analyzed. All the hematopoietic cell lineages were located within the mesenchyme and adjacent to pancreatic epithelial cells.

### Characterization of fetal pancreatic resident macrophages

Tissue-resident macrophages have a unique ontogeny since they can be generated in the yolk sac from primitive and erythro-myeloid progenitors or from definitive hematopoietic stem cells^15,78,79^. Macrophages derived from yolk sac hematopoiesis migrate into the developing embryo to ultimately reside in different tissue niches where they contribute to tissue homeostasis, native immunity, and organogenesis^80-82^. To gain insights into the fate of the pancreatic fetal macrophages, we compared the macrophage clusters from our dataset to those from a publicly available snRNAseq dataset of neonatal and adult human pancreas^14^. This comparison indicated that neonatal and adult macrophages do not express *CD36* and *LYVE1* (Figure 3A), and that, among the fetal macrophage populations, the perivascular macrophages (*CD36*+) have the lowest similarity to neonatal macrophages (Figure 3B), suggesting that *CD36*+ perivascular macrophages might be exclusive to the developing fetal pancreas. To validate the snRNAseq data, we performed immunostaining and CLSM analysis of pancreatic sections at 14, 16 and 18 PCW and confirmed the presence of CD36+ as well as LYVE1+ macrophages in fetal tissues (Figure 3C).

**Figure 3.**
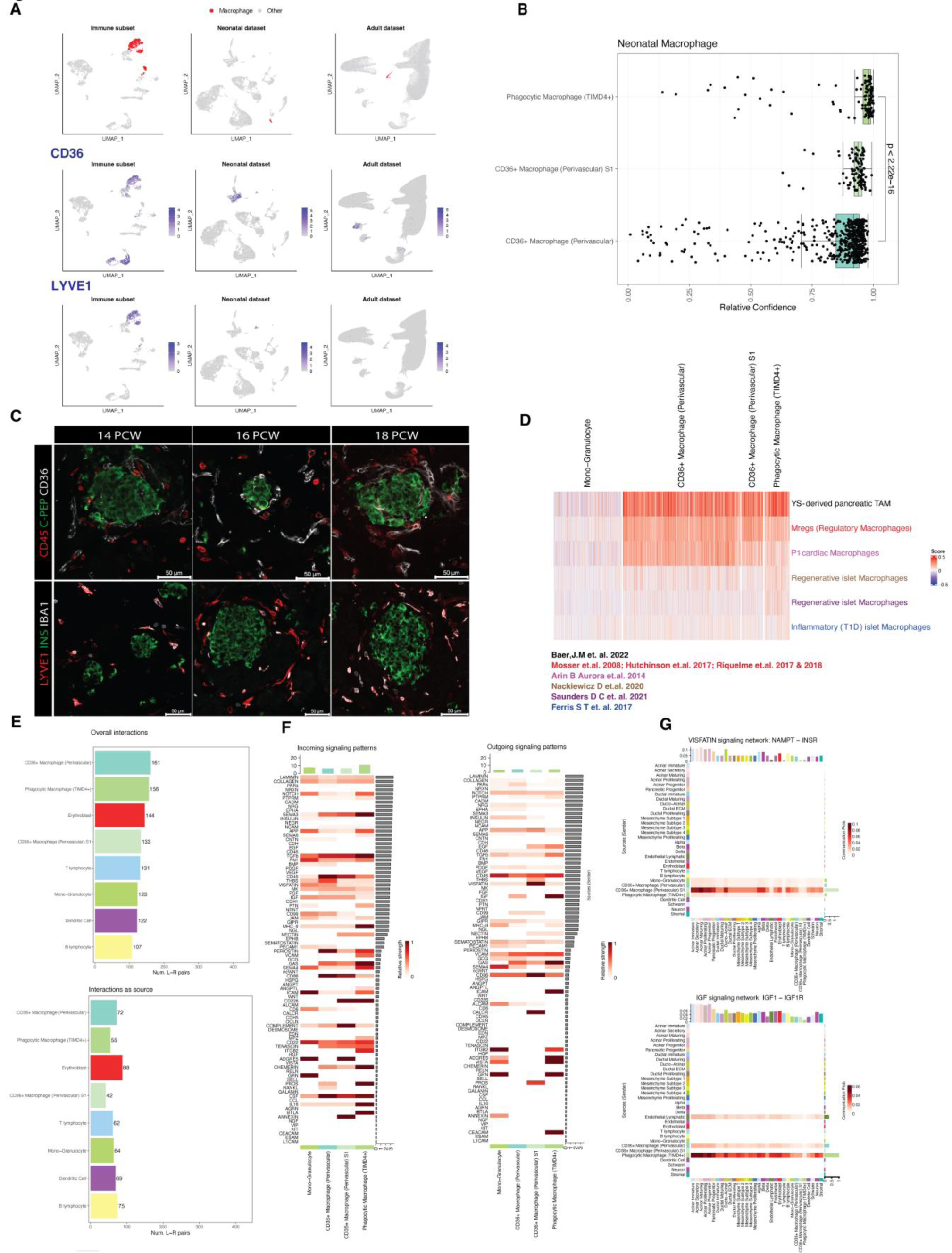
Heterogeneity of the fetal resident pancreatic macrophages. **A**, UMAP visualization of Macrophages clusters (red) identified in the fetal immune subset (this study), the neonatal and adult datasets (from Tosti et al., 2020). *CD36* and *LYVE* expression (blue) within the fetal immune subset (this study) compared to the neonatal, and adult pancreas datasets (from Tosti et al., 2020). **B**, Boxplot showing comparison by canonical correlation analysis between the fetal pancreatic macrophage subpopulations and neonatal pancreatic macrophages (dataset from Tosti et al., 2020). **C**, CLSM images of fetal tissue section at 14-18 PCW highlighting CD36+ macrophage (perivascular) and LYVE1+ macrophage (perivascular) within the developing pancreatic epithelium. **D**, Heatmaps of signature scores (rows) for bulk-derived expression of macrophages populations compared to fetal pancreatic macrophages from this study (columns). **E**, Bar plot of the number of overall interactions and interactions as source (outgoing interactions) detected by L-R analysis from Figure 1E, within the hematopoietic cells in the fetal pancreas. **F**, Heatmap summarizing outgoing and incoming signaling pathway interactions within the main macrophage subpopulations identified in Figure 2C. Top and left bar plots to depict total interaction strength of each of the macrophage populations and signaling pathways respectively across the dataset. **G**, Heatmap of predicted VISFATIN and IGF signaling pathway interactions by L-R analysis across all the major cell types in the fetal pancreas. The top bar plots depict the total interaction strength in each pathway by cell type.

Next, we interrogated previously reported macrophage RNA signatures (derived from independent transcriptome bulk studies) and performed correlation analyses to compare these datasets to the phenotype of fetal pancreatic macrophages from this study (Figure 3D; Table 1). While we found similarities with both YS-derived pancreatic Tumor Associated Macrophages (TAM)^83,84,85^ and Regulatory Macrophages (Mregs)^86-89^, fetal macrophages correlated poorly with regenerative islet macrophages^90,91^ and inflammatory (T1D) islet macrophages^92^ (Figure 3D). Remarkably, perivascular CD36+ macrophages showed high similarities to postnatal day 1 (P1) cardiac macrophages, which have been shown to have regenerative properties^74^ (Figure 3D; Figure S7C).

To investigate the heterogeneity and the molecular crosstalk between fetal macrophage subpopulations and pancreatic cells, we focused the L-R interaction analysis (from Figure 1E) on macrophages. When looking at the numbers of overall L-R interactions and L-R interaction as the source, macrophages scored the highest among the immune cells, emphasizing a key role played by these cells within the developmental pancreatic niche (Figure 3E).

We did not identify any quantitative differences in the number of L-R interactions between macrophage populations, but we observed that all these populations participated in numerous signaling pathways that were activated during pancreatic fetal development (Figure 3F). Noticeably, the *CD36+* perivascular macrophage S1 cluster is the major source of *VISFATIN*, also known as extracellular Nicotinamide phosphoribosyl transferase (e*NAMPT*) (Figure 3G). This is the rate-limiting NAD+ biosynthetic enzyme known to convert NAM into NMN, sustaining NAD synthesis and energetic metabolism^93-95^. Mouse studies have shown that physiological concentrations of eNAMPT regulate glucose-stimulated insulin secretion, and elevated concentrations of eNAMPT are found in T2D patients and are associated with beta cells dysfunction^96^. In contrast to mouse studies, where the mesenchyme is the major source of IGF signaling^97^ during pancreas development, here we show that phagocytic macrophages together with *CD36+* perivascular macrophages and endothelial lymphatic cells are the major source of the *IGF1* signalling network^90,98^ (Figure 3G). While IGF signaling does not play a role in beta cell development^99^, IGF1 has been shown to be necessary for beta cell function in rodents^100,101^. These findings uncover novel signaling cross-talks between macrophages and the pancreatic epithelium that may orchestrate pancreas development and native immunity.

### Macrophages-endocrine crosstalk during hESC differentiation towards endocrine cells

To investigate the role of *CD36+* perivascular macrophages during pancreatic development, we modelled hESC-derived macrophage/endocrine cell interactions *in vitro*. Using a protocol that recapitulates human yolk sac hematopoiesis^15^ (Figure 4A), we generated KDR+CD235+ hematopoietic mesoderm followed by CD43+ hematopoietic progenitors (Figure S8A). CD43+ cells were purified and differentiated further to derive macrophages (Figure 4B), referred to hereafter as embryonic macrophages (eMACs), that display expression of the surface marker of mature macrophages (CD45, CD64, CD11b, CD14, CD163) (Figure 4C) as well as expressing key markers of the fetal perivascular macrophages identified by this study (CD36 and MRC1) (Figure 4D). To study the role of eMACs during pancreatic development, we established a co-culture system that allowed us to monitor the formation of pancreatic progenitors (PP) and their commitment to endocrine cells in the presence of eMACs (Figure 4E). Whole mount staining and CLSM showed that the two cell types aggregated to form 3D structures consisting of an inner core of PDX1+/NKX6-1+ cells surrounded by a dense mantel of CD45+ macrophages (Figure 4F). Occasionally, some CD45+ cells were found in the core of the aggregates, intermingling with the developing endocrine islets cells (Figure S8B). This co-culture system did not affect the eMAC phenotype nor change the frequency, proliferation, or phenotype of PP cells (Figure S8C; Figure S9A-D). In contrast, prolonged co-culture of eMACs with developing endocrine cells showed a beneficial effect on the viability and frequency of the endocrine cells. Notably, eMAC-Endo organoids, composed of ∼40% CD45+ eMACs, yielded higher frequencies and numbers of Chromogranin A (CHGA)+/NKX6.1+ endocrine cells than endocrine aggregates alone (Figure 4G,H; Figure S9E-F).

**Figure 4.**
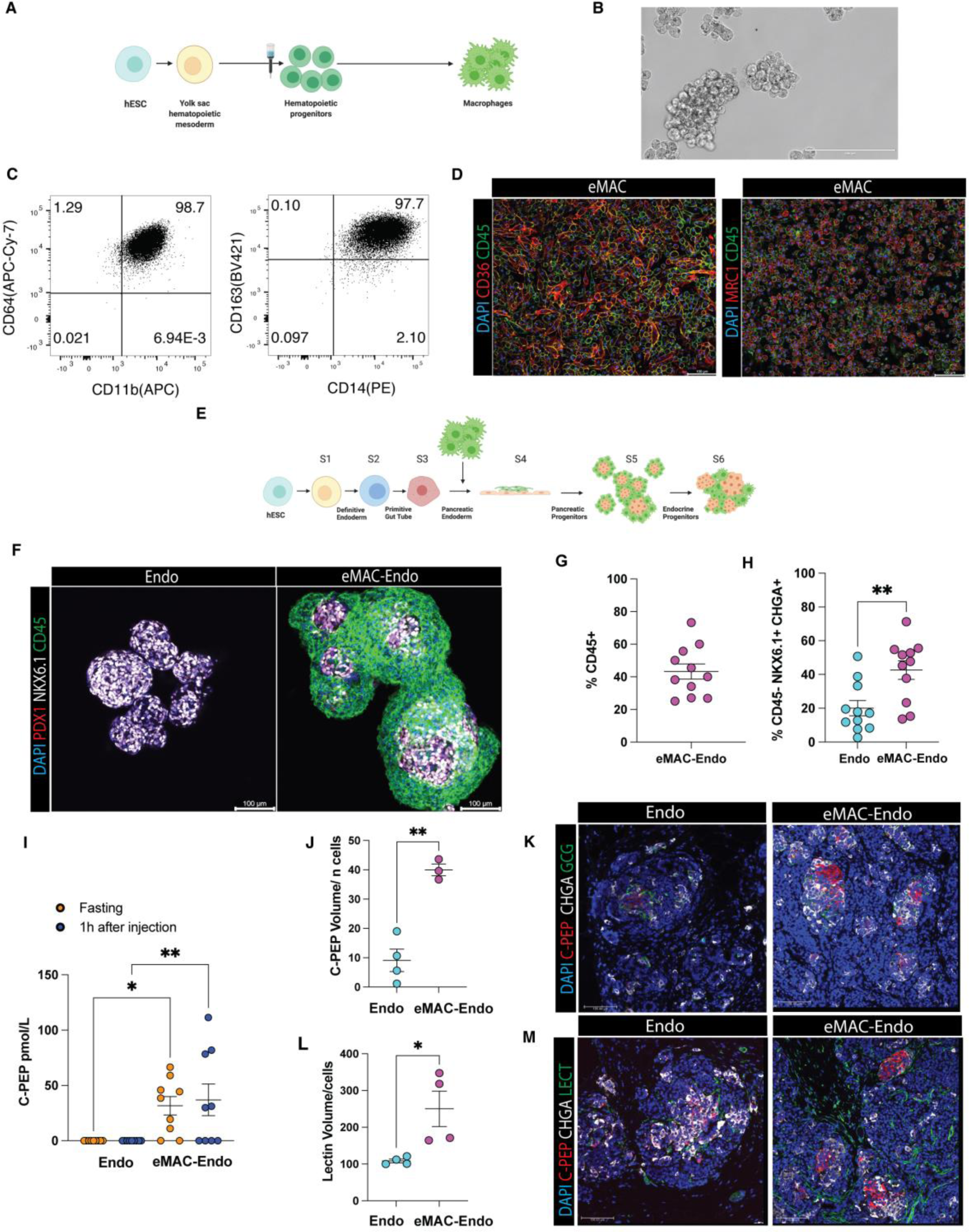
hESC-derived macrophages-endocrine coculture support endocrinogenesis in vitro and in vivo. **A**, Schematic of protocol to generate hESC-derived embryonic macrophages (eMAC). **B**, Representative bright field image of hESC-derived eMAC cells. **C**, Flow cytometry profile of CD64, CD11b, CD14 and CD163 in hESC-derived eMACs. **D**, CLSM images of hESC-derived eMACs expressing CD45, MRC1 and CD36. **E**, Schematic of hESC derived eMAC and developing endocrine cells co-culture (eMAC-Endo). **F**, CLSM images of hESC-derived endocrine organoids (Endo) and hESC-derived macrophages and endocrine organoids (eMAC-Endo) stained for PDX1 (red), NKX6-1 (white) and CD45 (green). **G**, Flow cytometry quantification of CD45+ cells within the eMAC-Endo organoids, (n=11 independent experiments, error bar represents SEM). **H**, Flow cytometry quantification of CD45-CHGA+NKX6.1+ endocrine cell generated without (Endo) and with (eMAC-Endo) macrophages (n=11 independent experiments, error bar represents SEM, unpaired t-test, **p value= 0.0047). **I**, Circulating human C-peptide was detected upon glucose-stimulated insulin secretion (GSIS) test in mice following 6-week post-subcutaneous transplantation with either Endo or eMAC-Endo cells (n=9 mice, 3 independent experiments, error bar represents SEM, one-way ANOVA, p-value: *=0.0218; **=0.007). **J,K**, Quantification of C-peptide+ volume (n=7 mice from 3 independent experiments, error bar represents SEM, unpaired t-test, **p value= 0.0014) and CLSM images of subcutaneous pancreatic graft from Endo and eMAC-Endo at 6 weeks post-transplant stained with c-peptide (red), GCG (green) and CHGA (white). **L,M** Quantification of Lectin+ volume (n=8 mice from 3 independent experiments, error bar represents SEM, unpaired t-test, *p value= 0.026) and CLSM images of representative subcutaneous pancreatic graft from Endo and eMAC-Endo at 6 weeks post-transplant stained with C-peptide (red), CHGA (white) and lectin (green).

To assess whether eMACs could provide a supportive environment for developing endocrine cells *in vivo*, we co-transplanted eMACs and endocrine aggregates (eMAC-Endo) in the subcutaneous space of normoglycemic immunocompromised mice and followed their engraftment and maturation. The subcutaneous space allows for ease of implantation, monitoring, and retrieval, making it an ideal site for transplantation, yet it brings some challenges in terms of vascular density, oxygen availability and mechanical stress^102,103^. Remarkably, 6-weeks post-transplantation, systemic human C-Peptide was detected only in mice transplanted with eMAC-Endo organoids (Figure 4I). Accordingly, CLSM analysis of the grafts at 6 weeks post-transplantation showed a greater frequency of mono-hormonal C-peptide + endocrine cells in the explants from eMAC-Endo aggregates compared to those from the Endo only group (Figure 4J,K). Furthermore, lectin immunostaining showed an extensive islet vascular architecture within grafts derived from eMAC-Endo compared to the Endo only grafts (Figure 4L,M). Taken together, these findings show that eMACs promote the survival and commitment of endocrine cells *in vitro* and improve engraftment and vascularization following transplantation *in vivo*. These data support a model where fetal macrophages may play a key role during pancreas development by promoting endocrinogenesis and angiogenesis and pave the way to the use of macrophages for the generation of transplantable hESC-derived endocrine organoids with therapeutic purposes for individuals with T1D.

## DISCUSSION

By combining snRNASeq and histological analysis, we explored and curated cell type annotation, ligand-receptor (L-R) pairing, and cell subtype-specific gene expression of the human fetal pancreas during islet development and morphogenesis. We detected cell diversity and heterogeneity among epithelial and non-epithelial cells and discovered novel transcriptional signatures amongst the major cell types and using computation tools, we deconstructed the signaling pathway network among pancreatic epithelial and niche cells. We generated a map of cell-cell interactions and biological pathways that coordinate pancreas-specific physiology and maturation during human fetal development, providing a reference on previously unknown signaling axes that contribute to pancreatic development. Focusing on the pancreatic hematopoietic niche, we created a blueprint of the composition and phenotype of this compartment during fetal development, which is a critical time for organ immune tolerance establishment. Understanding the composition and phenotype of the pancreatic hematopoietic niche during fetal development is crucial for gaining insights into the mechanisms underlying the development of autoimmune diseases like T1D in the pediatric population. We showed that the fetal pancreas of the second trimester harbored a heterogenous T cells population with a naïve as well as a memory phenotype. Memory T cells are a specialized type of immune cell that are cable of recognizing and responding rapidly and effectively to specific antigens upon subsequent exposure^104^. The presence of this T cells population within the fetal pancreas indicates that the developing immune system is already competent to respond to specific antigens at this early stage of fetal development.

Accordingly, we also detected immature pre-B cells and B cells residing in the human fetal pancreas of the second trimester This suggest that the developing immune system within the fetal pancreas, is already equipped to generate pancreatic autoantigens, potentially playing a role in the establishment of pancreas specific immune tolerance. Pancreatic tissue-resident NK and ILC3 cells were detected in our data set, suggesting an unexplored role of these cells in tissue homeostasis similar to previous reports from other GI organs^59^. Future comparison of these transcriptional signatures to datasets originating from donors with T1D and auto-antibody positive non-diabetic donors^105^ could ascertain the relevance of specific immune cell populations and their potential role in disease development.

Consistent with other studies reporting heterogeneity in time, phenotype, and location among macrophages, we identified two subpopulations of fetal pancreatic macrophages with distinct transcriptomic profiles and potential differences in developmental origins and functions: *LYVE1+CD36+* perivascular macrophages and *LYVE1-TIMD4+* phagocytic macrophages. While recent work highlights the co-expression of *TIMD4, LYVE1*, and *FOLR2* as core gene signatures of embryonic and fetal tissue-resident macrophages (TLF+ Macrophages) in humans^106^, our data show that the expression of *LYVE1* and *TIMD4* is mutually exclusive in the pancreatic fetal tissue-resident macrophages and *FOLR2* is not detected. It remains to be elucidated if this heterogeneity in macrophages is linked to their origin (primitive or definitive hematopoiesis), developmental stage or tissue geography (islet macrophages or acinar macrophages).

Intriguingly, the *LYVE1+CD36*+ macrophage population is not detected in human neonatal or adult pancreas, suggesting that they only exist during fetal time and share expression of *CD36* with endothelial and erythroid cells in the pancreas. Consistent with these data, the human fetal macrophages identified in our study are distinct from the monocyte-derived CCR2+ macrophage found in the neonatal mouse pancreas^13^.

These findings may suggest that different macrophage subsets are present at different developmental stages during pancreas development. The concept that macrophage populations are developmentally dynamic is not novel. For example, macrophages with regenerative capacity are only found at postnatal day 1 (P1) in the mouse heart^74^. Remarkably, *LYVE1+CD36+* perivascular macrophages from our study have high similarities to these regenerative P1 cardiac macrophages, suggestive of their role in angiogenesis and tissue remodelling.

Further, computational analysis indicated shared characteristics between fetal pancreatic macrophages (from this study) and YS-derived pancreatic Tumor Associated Macrophages (TAM), as well as human Regulatory Macrophages (Mregs). Mregs are immunosuppressive myeloid cells often found in solid tumors, where they contribute to creating an immunosuppressive environment and facilitating the tumor’s ability to evade the immune system. *In vitro*, human and mouse monocyte-derived macrophages can be induced to acquire an immunosuppressive and regulatory phenotype. These macrophages have been shown to provide temporary protection against type 1 diabetes in non-obese diabetic (NOD) mice^107^ and increase heart allograft survival in fully immunocompetent murine recipients^108^. This, together with our findings, highlight the crucial need to comprehend the physiological significance of macrophage heterogeneity and their potential therapeutic benefits for cell replacement therapy. In accordance with this theory, we showed that hESC-derived eMAC resembling *LYVE1+CD36+* perivascular macrophages promoted *in vitro* differentiation of hESC-derived endocrine cells and supported their engraftment *in vivo*. This is the first protocol establishing an efficient co-culture system between macrophages and pancreatic endocrine cells and demonstrating the beneficial effect of macrophages on endocrine development. The generation of macrophages-endocrine organoids clearly supports endocrinogenesis and islet-like-morphogenesis *in vitro*. Moreover, grafts originating from eMAC-endocrine organoids transplanted subcutaneously had increased vessel density compared to grafts originating from endocrine cells alone, suggesting that macrophages may directly or indirectly promote vascularization. Further studies will be required to clarify the molecular pathways guiding macrophage-endocrine crosstalk during endocrine differentiation and the mechanism supporting cell engraftment. Additionally, this co-culture system offers a novel platform for immunological studies targeting autoimmunity in T1D and could be also applied to other cell transplant strategies to facilitate engraftment in the subcutaneous space.

### Limitations of the study

While we were able to sequence a large number of cells, our study is limited by the number of samples and the developmental time analyzed, which is restricted to the first weeks of fetal pancreatic development. To further delineate the specificity and dynamics of the hematopoietic niche we described, it would be of great interest to complement our dataset with additional studies using embryonic samples. Additionally, transcriptomic data from snRNAseq may be limited in its ability to detect only highly abundant RNA transcripts. However, it offers the advantage of eliminating cell stress associated with single cell dissociation of tissue. In addition, lack of spatial transcriptome information regarding the hematopoietic fetal pancreatic niche might have hindered valuable insight in the heterogeneity of the macrophage subpopulations identified. In this study, we focused on unpolarized hESC derived primitive macrophages to assess their ability to support endocrinogenesis both *in vitro* and *in vivo*. However, exploring the role of polarized macrophages (M1 or M2) or macrophages from alternative sources (derived from cord blood or from monocyte) could further enhance our understanding of the potential contributions of these immune cells during endocrine development as well as their application for cell transplantation.

## Acknowledgements

The authors thank the donors, RCWIH Biobank, the Lunenfeld-Tanenbaum Research Institute, and the Mount Sinai Hospital/UHN Department of Obstetrics and Gynaecology for the human specimens used in this study (https://biobank.lunenfeld.ca). We thank Carolin Sebastiany for technical support with immunostainings and Amanda Oakie for helpful comments during manuscript preparation. This work was supported by a New Idea Grant from the Ontario Institute for Regenerative Medicine, a grant from the Howard Webster Foundation and the Toronto General and Western Hospital Foundation. A.M. was supported by an advanced post-doctoral fellowship from the Juvenile Diabetes Research Foundation (3-APF-2018-585). R.S. was supported by post-doctoral fellowships from the Juvenile Diabetes Research Foundation (1-PDF-2019-716 A-N) and (3-PDF-2020-954 A-N). AJG and CC were partially supported by CIHR project grant (PJT-180525) and used equipment supported by Canada Foundation for Innovation John R. Evans Leaders Fund (JELF). M.H. Atkins was supported by a Canadian Institutes of Health Research Banting and Best Doctoral Research Award.

## METHODS

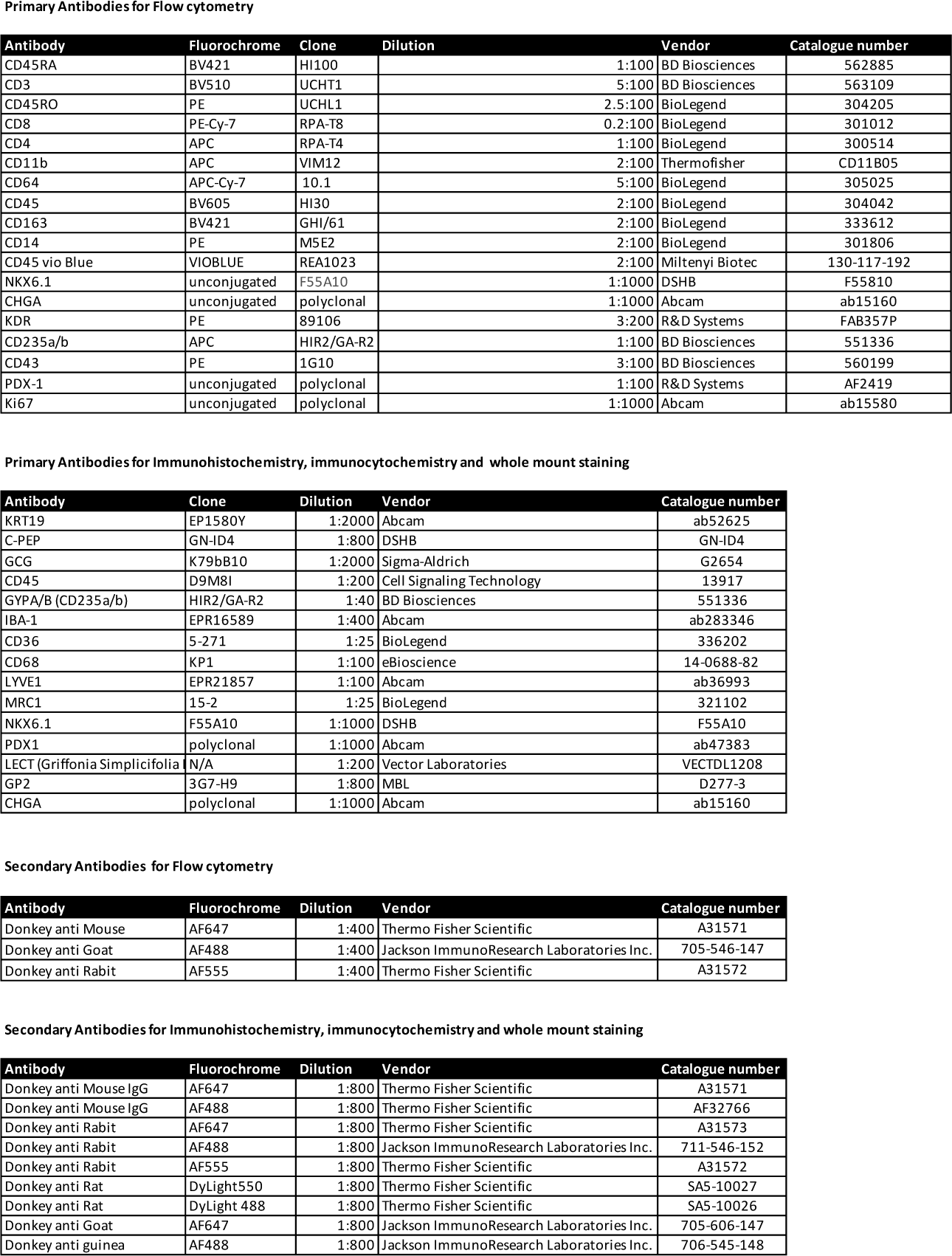
KEY RESOURCES TABLE.

## RESOURCE AVAILABILITY

### Lead contact

Further information and requests for reagents may be directed to, and will be fulfilled by the lead contacts, Dr. Maria Cristina Nostro and Dr Adriana Migliorini

### Materials availability

This study did not generate new unique reagents.

### Data and code availability

All SnNuc-Seq data have been deposited at GEO and will be publicly available as of the date of publication with accession number.

This paper does not report original code. All SnRNAseq related analyses in this study were performed using standard protocols and packages as previously described in the methods details sections. Code is available upon request to corresponding authors or to the laboratory of Dr. Faiyaz Notta.

## EXPERIMENTAL MODEL AND SUBJECT DETAILS

### Animals

The NOD. Cg-Prkdcscid Il2rgtm1Wjl/SzJ (NSG) male mice were purchase from Jackson Laboratories and bred in our facility at University Health Network, Toronto, Canada. All animal experiments described in this study were approved by the Institutional Animal Care & Use Committee (IACUC) of University Health Network.

### Human fetal pancreatic samples

Normal, de-identified human pancreatic fetal tissues were obtained from the Research Centre for Women’s and Infant’s Health (RCWIH) biobank (https://biobank.lunenfeld.ca/) with approval from the Research Ethics Board of the Mount Sinai Hospital (REB:18-0188-E) and University Health Network (UHN) Research Ethics Board (REB:17-6270). Samples were collected with written informed consent according to the procedures approved by the Research Ethics Board of the Mount Sinai Hospital. Fetal developmental stages assignment was performed by measuring the femur length, biparietal diameter, and foot length and compared to a standard growth chart^1^. Specimens were collected andol stored on ice in a sterile saline solution before microdissection and tissue processing for downstream analysis.

### Human embryonic stem cells

The human embryonic stem cell (hESC) line H1 (NIH code WA01) was obtained from WiCell Research Institute (Madison, WI). All experiments using hESCs were approved by the Stem Cell Oversight Committee (Canadian Institute of Health Research). The hESC H1 cell line tested negative for Mycoplasma contamination (Microbiology Lab, Sick Kids). Stem cells were maintained as a monolayer in hESC growth media on irradiated mouse embryonic feeder cells (iMEF’s) as previously described^2^ or in StemMACS™ iPS-Brew XF media (Miltenyi Biotec) on Geltrex (Gibco) coated 6-well plates (BD, Falcon) for feeder-free differentiation.

## METHOD DETAILS

### Differentiation of hESCs towards macrophages and pancreatic endocrine cells

Hematopoietic differentiation from hESCs (day 0) to CD43^+^ hematopoietic progenitors (day 9) was performed as previously described in^3^. On day 9 of differentiation, cells were dissociated with 0.25% trypsin-EDTA (Corning) for 5 minutes at 37°C before incubation in 0.2% collagenase type II (Worthington Biochemical) for 1 hour at 37°C. 1×10^8^ cells/mL were then stained with CD43 microbeads (diluted 1:5, Miltenyi Biotec) for 15 minutes at 4°C. The cells were sorted on an LS column (Miltenyi Biotec) and the bound fraction (CD43^+^ cells) was collected and further cultured to generate macrophages. Sorted CD43^+^ cells were transferred to diluted StemPro-34 media (Gibco) supplemented with 100 ng/mL SCF (R&D Systems), 50 ng/mL IL-3 (R&D Systems), and 30 ng/mL M-CSF (R&D Systems) and cultured in suspension for 3 days at a density of 1×10^5^ cells/mL in normoxia condition (37°C, 5% CO2). Thereafter, the cells were maintained in StemPro-34 media (Gibco) diluted 1:4 in IMDM and supplemented with 30 ng/mL M-CSF (R&D Systems). Cells were passaged as needed, based on culture density. The media was changed every 3 days for a minimum of 12 days before their use in co-culture studies. Flow cytometry analysis was used to assess the efficiency of differentiation toward macrophages.

Pancreatic endocrine differentiation from hESCs cultured on irradiated mouse embryonic feeders (iMEFs) (day 0) to endocrine cells (day 20) was performed using the six-stages protocol (S0-S6) described in^4^ with the following modifications. At the end of S4, cells were enzymatically dissociated using TrypLE (Gibco) and plated at a density of 1-1.5×10^6^ cells/mL on polyheme (Poly(2-hydroxyethyl methacrylate), Sigma) coated tissue culture plates.

Pancreatic endocrine differentiation from feeder-free hESCs (day 0) to endocrine cells (day 20) was performed using the following six-stages protocol (S0-S6) adapted from previously published protocols^6-7^. Differentiation began when hESCs were approximately 90% confluent. All media were supplemented with 1% glutamine (Hyclone) and 1% Penicillin-Streptomycin (Hyclone).

S1 (Day 0) media consisted of MCDB131 (Wisent), 5mM D-(+)-glucose (Sigma), 1.5g/L NaHCO3 (Gibco), 0.5% Fatty Acid-free BSA (Proliant Biologicals), 100ng/ml Activin A (R&D Systems) and 3µM CHIR990210 (Tocris). S1 (Day 1) media consisted of MCDB 131, 5mM D-(+)-glucose, 1.5g/L NaHCO3, 0.5% Fatty Acid free BSA, 100ng/ml Activin A and 0.3µM CHIR990210. S1 (Day 2) media consisted of MCDB131, 5mM D-(+)-glucose, 1.5g/L NaHCO3, 0.5% Fatty Acid free BSA, and 100ng/ml Activin A (R&D Systems). S2 media (Day 3, 5) consisted of MCDB131, 5mM D-(+)-glucose, 1.5g/L NaHCO3, 0.5% Fatty Acid-free BSA, 50µg/mL L-Ascorbic acid (Sigma), 50ng/mL hFGF10 (R&D System) and 0.75µM Dorsomorphin (Sigma). S3 media (Day 6, 7) consisted of DMEM (high glucose, Gibco) supplemented with 1% vol/vol NeuroBrew-21 without Vitamin A (Miltenyi Biotec), 50µg/mL ascorbic acid, 50ng/mL hNoggin (R&D System), 50ng/mL hFGF10, 0.25µM SANT-1 (Tocris) and 2µM all-trans retinoic acid (RA) (Sigma). S4 media (Day 8, 10, 12) consisted of DMEM containing 1% vol/vol NeuroBrew-21 without Vitamin A, 50µg/mL L-Ascorbic acid, 50ng/mL hNoggin, 100ng/mL hEGF (R&D System), 9µM WIKI4 (Tocris) and 30 ng/mL MCSF (R&D Systems). At Day 13, cells were enzymatically dissociated using TrypLE and plated at a density of 1-1.5×10^6^ cells/ml on polyheme (Poly(2-hydroxyethyl methacrylate), Sigma) coated tissue culture plates in S5 media. S5 media (Day 13) consisted of MCDB131 supplemented with 1% vol/vol NeuroBrew-21 without Vitamin A, 1µM T3 (3,3’, 5-Triiiodo-L-thyronine, Sigma), 1.5g/L NaHCO3, 15mM D-(+)-glucose, 10µg/mL Heparin (Sigma), 0.25µM SANT-1, 10µM RepSox (Tocris), 100nM LDN193189 (Tocris), 10µM ZnSO4 (Sigma), 0.05µM all-trans RA,10µM Y27632 (Tocris), 10nl/mL DNase 1 bovine pancreas (MilliporeSigma) and 30 ng/mL M-CSF (R&D Systems). S6 media (Day 16, 20) consisted of MCDB131 supplemented with 1% vol/vol NeuroBrew-21 without Vitamin A, 1µM T3, 1.5g/L NaHCO3, 10µg/mL Heparin, 10µM RepSox, 100nM LDN193189, 10µM ZnSO4, 100nM DBZ (Tocris) and 10 ng/mL M-CSF.

### Generation and culture Endo-MAC organoids

hESCs were parallelly differentiated into macrophages and pancreatic endoderm as described above. At the beginning of S4 of the endocrine differentiation, fully differentiated macrophages were added to the pancreatic cell monolayer culture. The differentiation medium was supplemented with 30 ng/mL M-CSF for the first 8 days of co-culture and 10ng/ml thereafter. Endocrine organoids without macrophages (Endo) were also cultured with matching amounts of M-CSF. The number of macrophages added to S4 pancreatic cells was normalized to 5% of the total number of pancreatic cells. After 48 hours (h), most of the macrophages were attached to the epithelium and an additional 1 ml of S4 media was added to each well. The following day, a full media change was performed and differentiation toward endocrine progenitors was carried on as described above and supplemented with M-CSF.

### Nuclei isolation, 10x Genomics Library preparation, sequencing, and alignment

All tissues were processed by the Princess Margaret Genomics Centre (PMGC), UHN, Toronto (https://www.pmgenomics.ca/pmgenomics/). Briefly, 30-50mg of snap-frozen pancreatic tissue was cut in 1-2 mm^3^ on dry ice and transferred to cold lysis buffer (0.32mM sucrose, 5mM CaCl2, 3mM Mg (Ac)2, 20mM Tris-HCl, 0,1% Triton X-100, 0.1mM EDTA, and RNase Inhibitor (40U/ml)). The tissue was chopped and homogenized with strokes using a Dounce homogenizer until intact nuclei were visible upon SYBR Green Nucleic acid staining (Thermo Fischer Scientific). Homogenate was centrifuged for 10 minutes at 800xg at 4°C. Supernatant was carefully removed, and nuclei were resuspended in 2ml of wash/ resuspension buffer (PBS + 10% BSA, 0.2U/μl RNase Inhibitor) and centrifuged for 10 min at 800xg at 4°C. Nuclei were resuspended again in 1ml of wash/resuspension buffer, filtered using a 40µm Flow micell strainer (Sigma-Aldrich) and transferred to a 1.5ml LowBind tube on ice. Nuclei were stained with SYBR GREEN again and counted on a disposable Hemocytometer. Nuclei were diluted to the desired concentration and stained with DAPI for fluorescence-activated cell sorting (FACS, Aria Fusion cell sorter). The gating was performed on DAPI+ nuclei to exclude debris and damaged nuclei.

Sorted nuclei were used as input into the 10X Genomics single-cell 3’ v3.1 assay and processed as outlined by the user guide (https://support.10xgenomics.com/single-cell-gene-expression/library-prep/doc/user-guide-chromium-single-cell-3-reagent-kits-user-guide-v31-chemistry). Briefly, following counting, the appropriate volume for each sample was calculated for a target capture of 1×10^4^ nuclei. After droplet generation, samples were transferred onto a pre-chilled 96-well plate (Eppendorf), heat sealed and incubated overnight in a Veriti 96-well thermocycler PCR machine (Thermo Fisher). The next day, sample cDNA was recovered using Recovery Agent provided by 10x Genomics facility, and subsequently cleaned up using a Silane DynaBead (Thermo Fisher) mix as outlined by the user guide. Purified cDNA was amplified for 11 cycles before being cleaned up using SPRIselect beads (Beckman). Samples were diluted 4:1 elution buffer (Qiagen): cDNA and run on a Bioanalyzer (Agilent Technologies) to determine cDNA concentration. cDNA libraries were prepared as outlined by the Single Cell 3’ Reagent Kits v3.1 user guide with modifications to the PCR cycles based on the calculated cDNA concentration. The molarity of each library was calculated based on library size as measured using a bioanalyzer and qPCR amplification data (Roche). Samples were pooled and normalized to 1.5nM. The library pool was denatured using 0.2N NaOH (Sigma) for 8 minutes at room temperature and neutralized with 400mM Tris-HCL (Sigma). Library pools at a final concentration of 300pM were loaded to sequence on Novaseq 6000 (Illumina). Samples were sequenced with the following run parameters: Read 1-28 cycles, Read 2-90, and index 1-8 cycles. Droplet-based (10x) sequencing data were processed using CellRanger (v3.1, 10X Genomics). The data were aligned and quantified using the official CellRanger human reference genome GRCh38 3.0.0, with transcript *INS-IGF2* removed from the human GTF file to unmask the *INS* transcripts. Barcodes with a low expression representing non-cell-associated barcodes were removed as default within CellRanger.

### Quality control, normalization, clustering, and dimensionality reduction

Further processing of data was performed within the Seurat (v3.2.3) R package. Cells with high mitochondrial content (>3%) were removed. Doublets were removed using the approach described in^8^. Briefly, the relationship between UMI and gene count was fit to a smoothing curve and cells, not within 4 standard deviations, were removed.

The filtered data were normalized using SCTransform in Seurat as described by^9^. The top 3000 variable genes were also identified using SCTransform. Principal component analysis (PCA) on these variable genes was performed using RunPCA, and a shared nearest neighbour graph (k=50) was generated with 40 principal components using FindNeighbors. This was followed by Leiden clustering (FindClusters, resolution 0.8) and UMAP dimensionality reduction (RunUMAP, 50 neighbours). After this initial processing, a cluster of cells dominated by mitochondrial gene expression was removed. Normalization, clustering, and dimensionality reduction were subsequently repeated on this updated dataset, and the updated processing results were used for downstream analysis. This analysis was also applied to the immune and endocrine data subsets.

### Differential expression analysis, gene set enrichment analysis and annotation

Differential expression analysis was performed in Seurat using FindAllMarkers with default parameters (Wilcoxon rank sum test, two-sided, log fold change > 0.25, expression in > 10% of each compared cell population). The top 50 significantly differentially expressed genes per cluster (Bonferroni-corrected p-value < 0.01) were investigated and aligned with marker genes identified through a literature search to annotate each cluster. The acinar, ductal and mesenchyme clusters were further processed and analyzed to refine cluster annotations. Clusters identified as acinar cells were subsetted and the differential expression analysis using FindAllMarkers was re-run (Wilcoxon rank sum test, two-sided, log fold change > 0.1, expression in > 10% of each compared cell population). Differentially expressed genes (Bonferroni-corrected p-value < 0.1) were ranked by log fold change and used for gene set enrichment analysis (GSEA) using the fgsea R package. MSigDB (Broad Institute) Hallmark gene sets and GO gene sets were used for GSEA. This analysis was also performed on ductal and mesenchyme cell clusters.

### Cell cycle scoring and pseudotime analysis

Single cells were scored for S phase genes and G2/M phase genes, and then designated a phase of S, G2/M, or G1 using CellCycleScoring in Seurat. Pseudotime and trajectory analyses were performed on the endocrine clusters using Monocle3 (v0.2.3). Dimensionality reduction and clustering were re-runs within Monocle3 on the endocrine subset with parameters closely matching those used in Seurat. The trajectory graph was built using learn_graph (default parameters), and the root of the Monocle3 trajectory graph was determined by high SOX4 expression. Genes showing the variation over pseudotime were identified using Moran’s I statistic with graph_test on the principal graph. The false discovery rate cut-off was set at 0.05. This analysis was performed for each specific cell lineage. The trajectory graph from the Monocle3 analysis was visualized using the tree graph layout in igraph (v1.2.6), and other pseudotime data were transferred to Seurat for further visualization.

### Ligand-receptor analysis

The dataset, featuring all cell type clusters, was used for ligand-receptor analysis. Ligand-receptor analysis was performed using CellChat (v1.1.0, default parameters) with the CellChat human ligand-receptor dataset.

### Comparison with neonatal macrophage dataset

The neonatal pancreas dataset from Tosti *et al.,*^10^ was obtained as a pre-processed Seurat object. The original cell type annotations of this dataset were used as a reference to perform Seurat’s Canonical Correlation Analysis (CCA)^11^ of the fetal immune subset. Both CCA functions, FindTransferAnchors and TransferData, were run using 50 principal components to produce confidence scores matching the neonatal cell types to each cell found in the fetal immune subset.

### Literature gene set scoring of macrophages

Gene sets from the literature were scored in all macrophage cells using Seurat’s AddModuleScore (default parameters). Briefly, the score is calculated as the average expression of the genes in the set subtracted by the average expression of reference genes randomly selected from the matching expression bin. For gene sets collected from mouse data, mouse genes were converted to human genes using one-to-one orthologues. Mouse genes without one-to-one orthologues were omitted.

### Immunohistochemistry, immunocytochemistry, whole mount staining and laser confocal microscopy

Fetal pancreatic specimens were fixed in 4% PFA overnight at 4°C and incubated in sucrose gradients before embedding in OCT and storage at −80°C. Tissue frozen sections were defrosted, rehydrated in PBS, and washed with PBS-T (PBS + 0.1% Tween-20, Sigma). Permeabilization was performed for 15-20 minutes in PBS + 0.5% Triton X, followed by 3 washes in PBS-T. Sections were incubated for 2 hours at 4°C with blocking solution (PBS-T containing 10% FBS, 3% donkey serum (JacksonLab), and 0.3% BSA) before overnight staining with primary antibodies diluted in blocking solution. Sections were washed the following day in PBS-T and stained overnight at 4°C with appropriate secondary antibodies diluted in a blocking solution. Nuclei were stained with DAPI diluted in PBS-T for 20 minutes followed by 3 washes in PBS-T. Sections were mounted with a coverslip using DAKO mounting solution (Thermo Fisher). µ-Slide 8 Well Glass Bottom (ibidi) was used to stain and image hESC-derived macrophages. The protocol used for staining macrophages was similar to the one described above with the following modifications: cells were fixed in 4% PFA for 20 minutes at room temperature, followed by washes in PBS-T and permeabilization for 10 minutes using PBS+0.5% Triton X (Sigma).

Whole-mount staining was performed on hESC-derived organoids following fixation in 4% PFA for 24 hours at 4°C. Organoids were first washed with a solution containing PBS, 0.1% gelatin, and 0.5% Triton X and permeabilized overnight using a solution containing PBS, 0.1% gelatin, 0.5% Triton X, and 1% saponin. Incubation for 24 hours with blocking solution (PBS-T containing 10% FBS, 3% donkey serum, and 0.3% BSA) was performed before antibody staining. Organoids were incubated on a shaker for 48-72 hours with primary antibodies diluted in PBS supplemented with 0.1% gelatin, 0.5% Triton X, 1% saponin, 5% FBS, 1.5% donkey serum, and 0.15% BSA. After 3 washes of 1 hour each with a solution containing PBS, 0.1% gelatin, 0.5% Triton X, and 1% saponin, organoids were incubated for 48 hours with secondary antibodies diluted in a solution of PBS supplemented with 0.1% gelatin, 0.5% Triton X, 1% saponin, 5% FBS, 1.5% donkey serum, and 0.15% BSA. After 3 washes of 2 hours each with a solution containing PBS, 0.1% gelatin, 0.5% Triton X, and 1% saponin, organoids were stained for 1 hour with DAPI diluted in PBS, 0.1% gelatin, 0.5% Triton X, and 1% saponin. Organoids were washed before mounting them on the coverslip using DAKO mounting solution (Thermo Fisher) and Secure-Seal spacer (Thermo Fisher).

Grafts explanted from the subcutaneous space were fixed in 10% formalin for 72 hours, followed by incubation in 70% ethanol and paraffin embedding. Sections were deparaffinized according to standard protocols, and heat-induced epitope retrieval was performed in TEG or citrate buffer. Sections were stained using the same protocol described above for frozen tissue sections. Details and dilutions of primary and secondary antibodies used are listed in the resource table.

Images of tissue sections and cells were captured using the Leica SP8 confocal microscope at the Advanced Optical Microscopy Facility, UHN. Brightness, contrast, and color adjustments of immunofluorescent staining images were made using Leica Application Suite X. Image quantification was performed using microscopy image analysis software Imaris 9.9.0.

### Flow cytometry

Cell monolayers and cell organoids were dissociated into single cells using TrypLE Express Enzyme (Gibco) at 37°C for 3-5 minutes (monolayer) or 12-15 minutes (organoids). The cells were washed and pelleted in PBS with 5% FBS and 10 µl/ml DNase. For surface marker staining, live cells were incubated with primary antibody and Fc-Blocking reagent (Miltenyi Biotec) in PBS with 5% FBS and 10 ng/ml DNase for 20 minutes at 4°C. Samples were washed twice and stained with DAPI before flow cytometry analyses. For surface markers staining of the fetal pancreas, the tissue was injected with 0.1% Collagenase and incubated for 30 minutes at 37°C. Samples were vortexed every 10 minutes. Upon complete tissue dissociation, cells were filtered, washed, and pelleted in PBS with 5% FBS and 10 ng/ml DNase. Single cells were stained with viability dye eFluor 520 (eBioscience) in PBS for 10 minutes before staining with Fluorophore-conjugated antibodies as described above. Cells were fixed in 1% PFA in preparation for flow cytometry analysis performed on a BD FACSymphony™ Cell Analyzer and BD LSRFortessa.

For intracellular marker staining, cells were fixed in BD Cytofix/Cytoperm solution (BD Biosciences) for 30 minutes, followed by incubation with primary antibodies overnight at 4°C diluted in Perm/Wash buffer (BD Biosciences). Next, samples were washed and incubated in secondary antibodies diluted in Perm/Wash buffer for 30 minutes at room temperature. Cells were washed and stored in PBS with 5% FBS and 10 ng/ml DNase in preparation for flow cytometry analyses. The details and dilutions of primary and secondary antibodies used are listed in the resource table. Flow cytometry data were analyzed using FlowJo software (BD Biosciences).

### Transplantation studies

Male NOD. Cg-Prkdcscid Il2rgtm1Wjl/SzJ (NSG) mice were acquired from Jackson Laboratories and bred in our facility at University Health Network, Toronto, Canada. Mice were treated in accordance with protocols approved by University Health Network. On the day of the transplant, Endo organoids and Mac-Endo organoids containing ∼2 x10^6^ S6 (D20) cells were embedded in hydrogel as previously described^12^ and implanted subcutaneously in 12-20-weeks old NSG male mice. Briefly, mice were anesthetized using 2% isoflurane. A left-flank incision was used to expose the skin after shaving and sterilization with isopropyl alcohol. Organoids were implanted beneath mouse skin using forceps. An *in vivo* Glucose Stimulated C-Peptide Secretion assay was performed at 6 weeks post-transplantation. Mice were fasted for 12 hours, and blood was collected at 0 and 60 min after glucose injection (3g/Kg of body weight). Human C-Peptide content in sera was quantified using the Ultrasensitive C-peptide ELISA kit (Mercodia). Mice were sacrificed at 6 weeks post-transplantation and grafts were collected for downstream analysis.

### Statistical analysis

GraphPad Prism 9 was used to perform statistical analysis of data not related to the snRNAseq. Results are expressed as standard error of the mean (± SEM) and details regarding the statistical test used are indicated in figure legends. Statistical analysis of snRNAseq was performed using R software and the Wilcoxon rank sum test, two-sided, was used to compare the confidence scores between the fetal macrophage populations reported in Figure.3B

## Supplementary Figures

**Figure S1.**
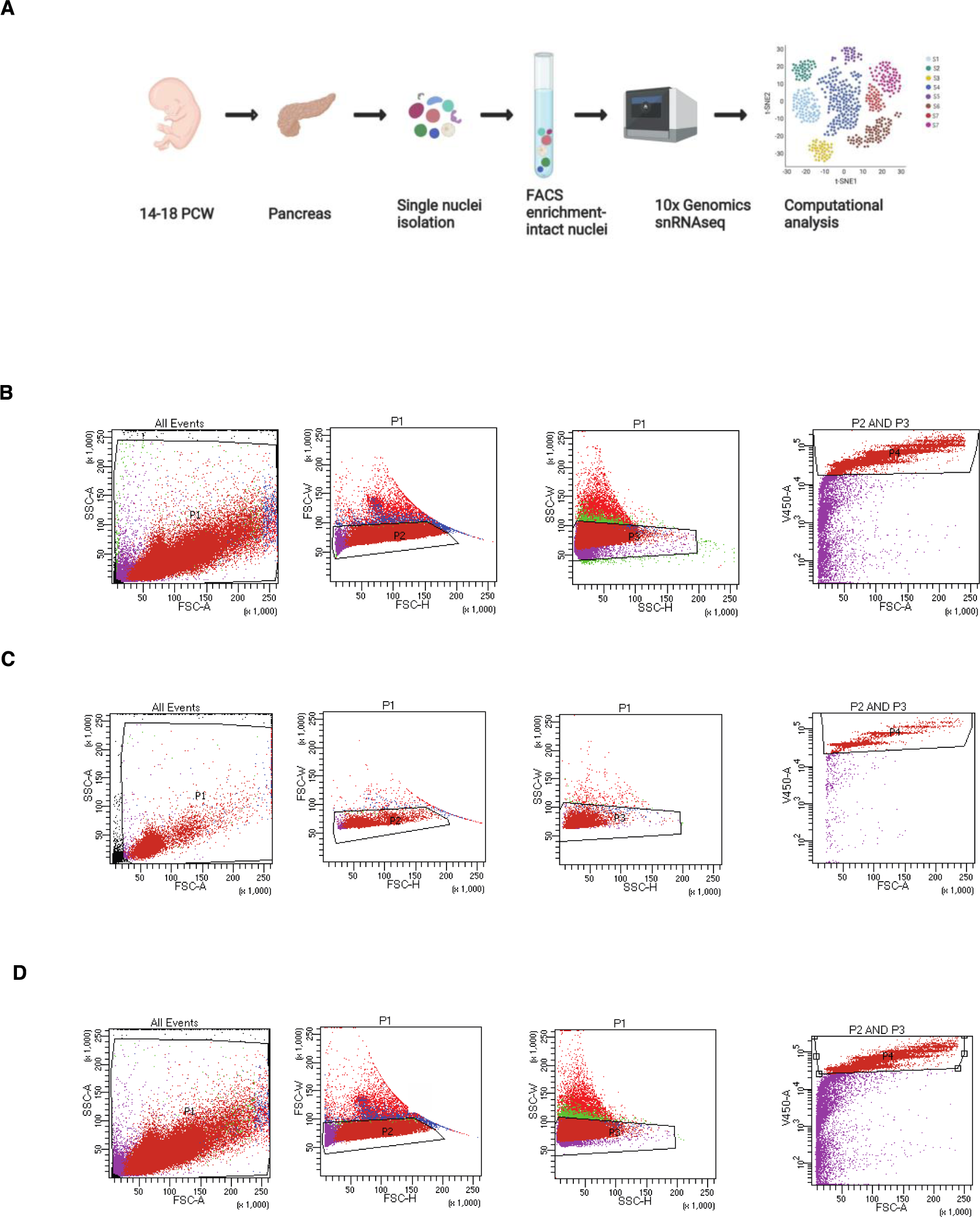
**A**,Schematic view of the experimental protocol for snRNAseq of the human fetal pancreas. **B-D** Representative Fluorescence Activated Cell Sorting (FACS) gating strategy for intact nuclei enrichment for snRNAseq for pancreatic fetal samples at 14, 16 and 18 PCW. All events were gated using forward and side scatter (P1), doublets were excluded by gating forward scatter high vs width (P2) and side scatter height and width (P3). Intact nuclei were sorted by excluding the V450-A (DAPI) negative cells within the P4 gate (P1+ P2 and P3)

**Figure S2.**
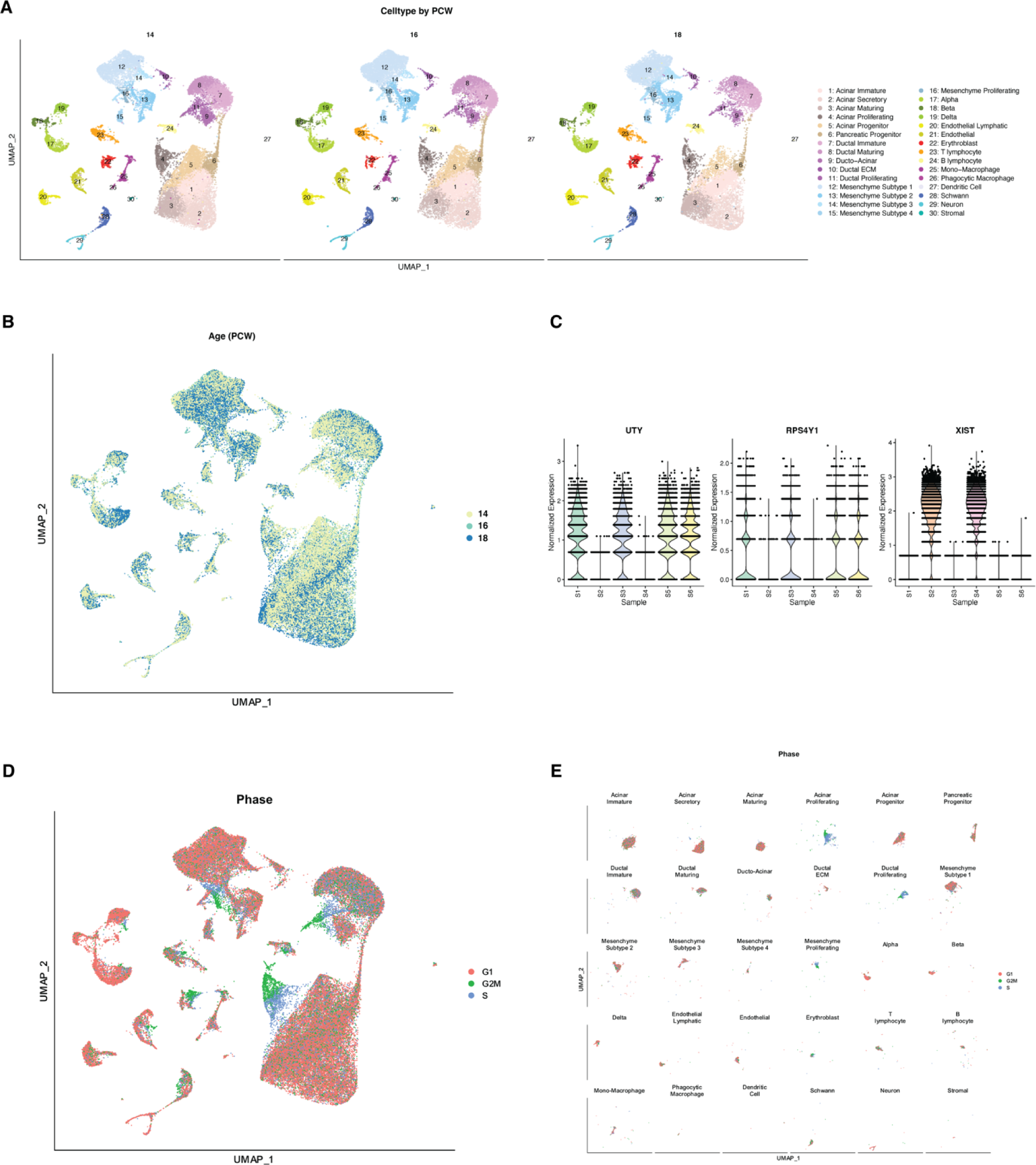
**A**,UMAP visualization of fetal pancreas cells grouped by developmental stages (n=6, 2 biologically independent samples per developmental time). **B**, UMAP visualization of fetal pancreas cells colored by developmental stage. **C**, Violin plots showing log-normalized expression in each sample of UTY, RPS4Y1 and XIST, which are upregulated in male and female samples, respectively. **D**, UMAP visualization for predicted cell cycle states of fetal pancreas cells. Colors indicate different phases of the cell cycle. **E**, UMAP visualization of cell cycle analysis of fetal pancreatic cells grouped by cell clusters.

**Figure S3.**
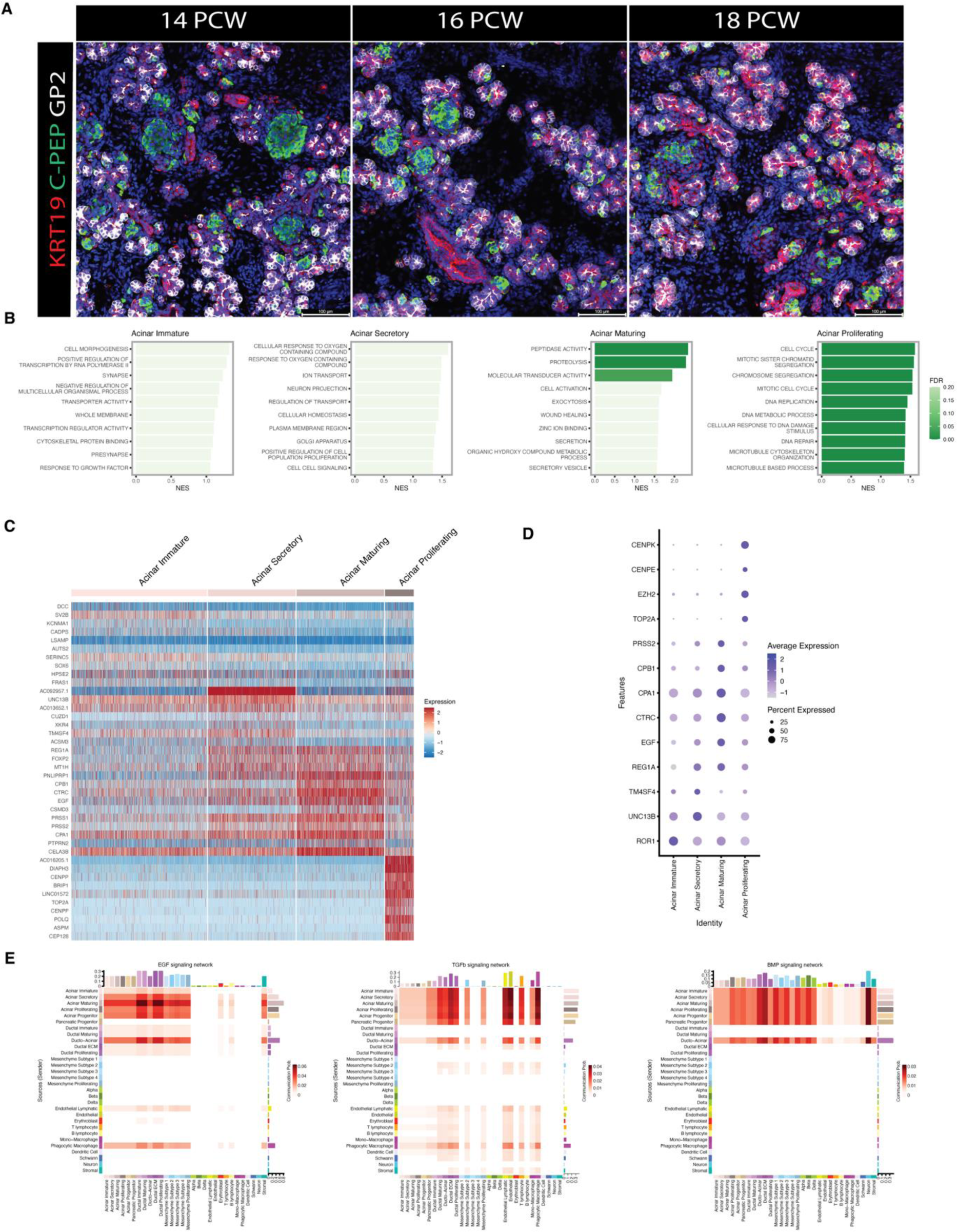
**A**,CLSM images of fetal pancreas stained for KRT19 (red), C-PEP (green) and GP2 (white) across three developmental stages (14-16-18 PCW). **B**, Bar plot of enriched gene programs among acinar cell populations detected by Gene Set Enrichment Analysis (GSEA). **C**, Heatmap of differentially expressed genes among acinar populations. **D**, Dotplot showing markers gene expression of annotated acinar cell populations**. E**, Heatmap of EGF, TGFb and BMP signaling pathway interactions predicted by L-R analysis (from Fig.1e) across all the major cell types in the fetal pancreas.

**Figure S4.**
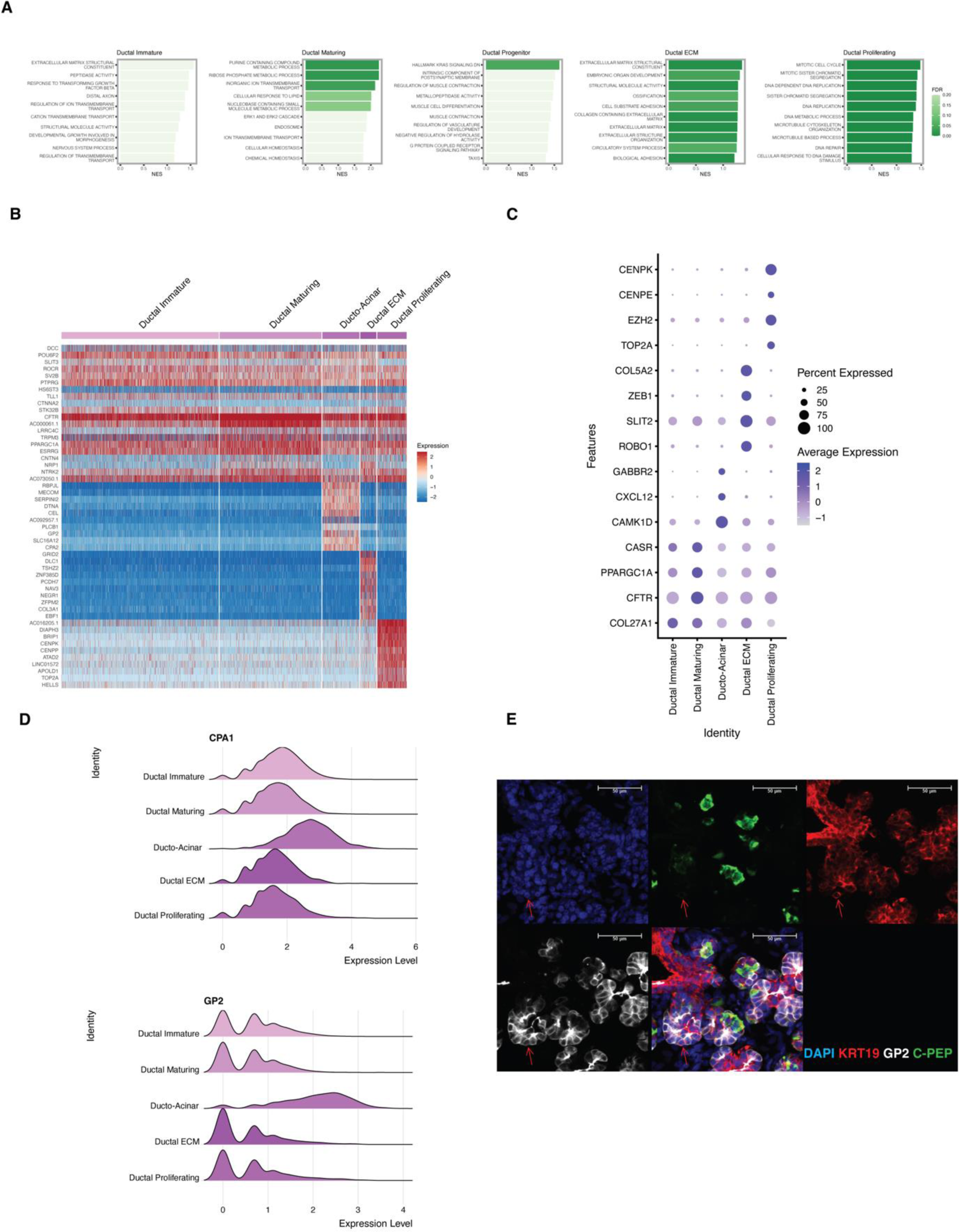
**A**,Bar plot of enriched gene programs among ductal cell populations detected by GSEA. **B**, Heatmap of differentially expressed genes among ductal populations. **C**, Dot plot showing marker gene expression of annotated ductal cell populations**. D**, Gene gradient expression of the acinar marker GP2 and CPA1 among ductal populations. **E**, CLSM image of fetal pancreatic epithelium stained for C-peptide (green), GP2 (white) and KRT19 (red) highlighting GP2+KRT19+ ducto-acinar cells.

**Figure S5.**
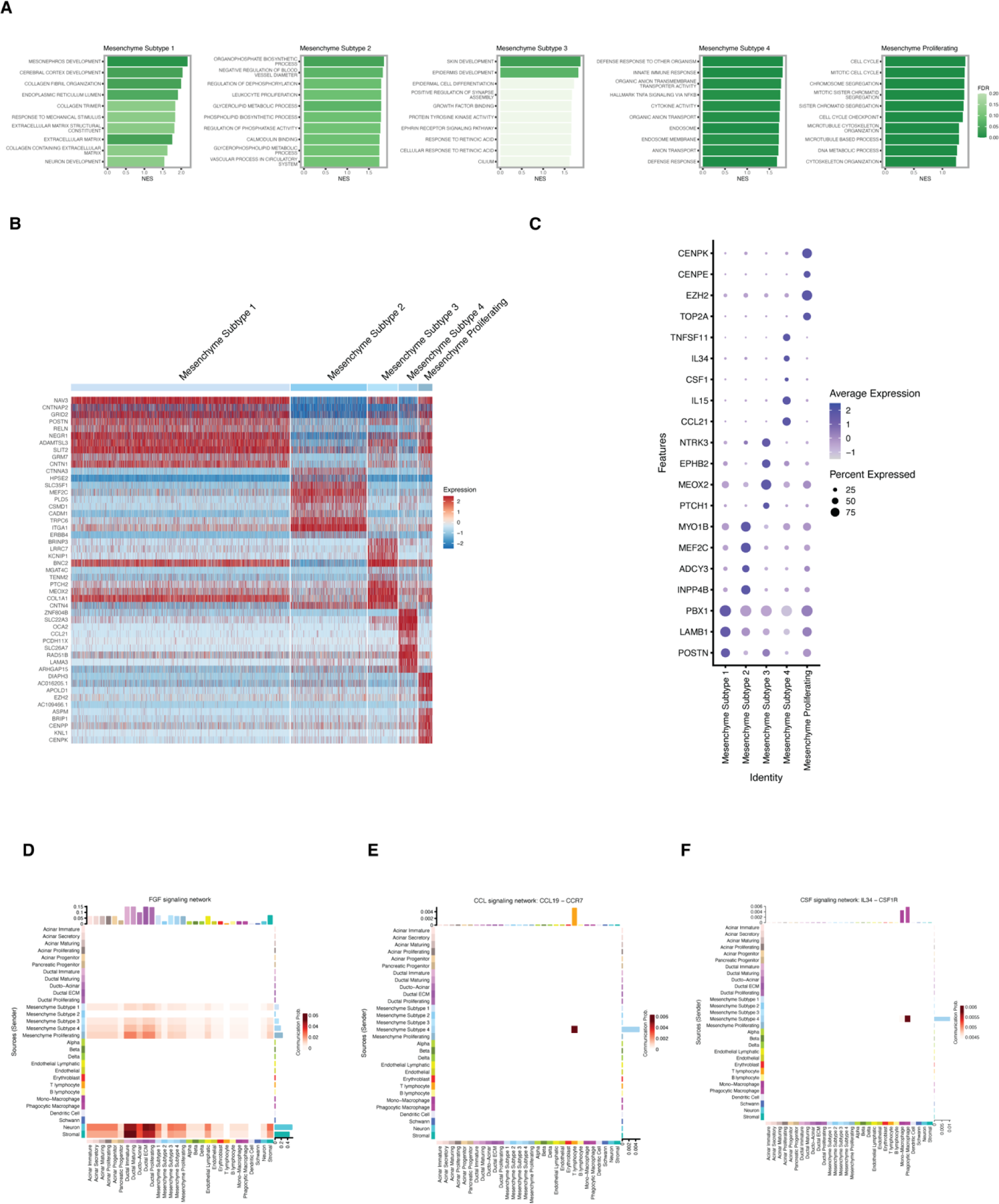
**A**,Bar plot of enriched gene programs among mesenchyme cell populations by GSEA. **B**, Heatmap of differentially expressed genes among mesenchyme subtype cell populations. **C**, Dot plot showing marker gene expression of annotated mesenchyme subtype cell populations**. D-F**, Heatmap of predicted FGF, CCL and CSF signaling pathway interactions by L-R analysis (from Fig. 1E) across all the major cell types in the fetal pancreas.

**Figure S6.**
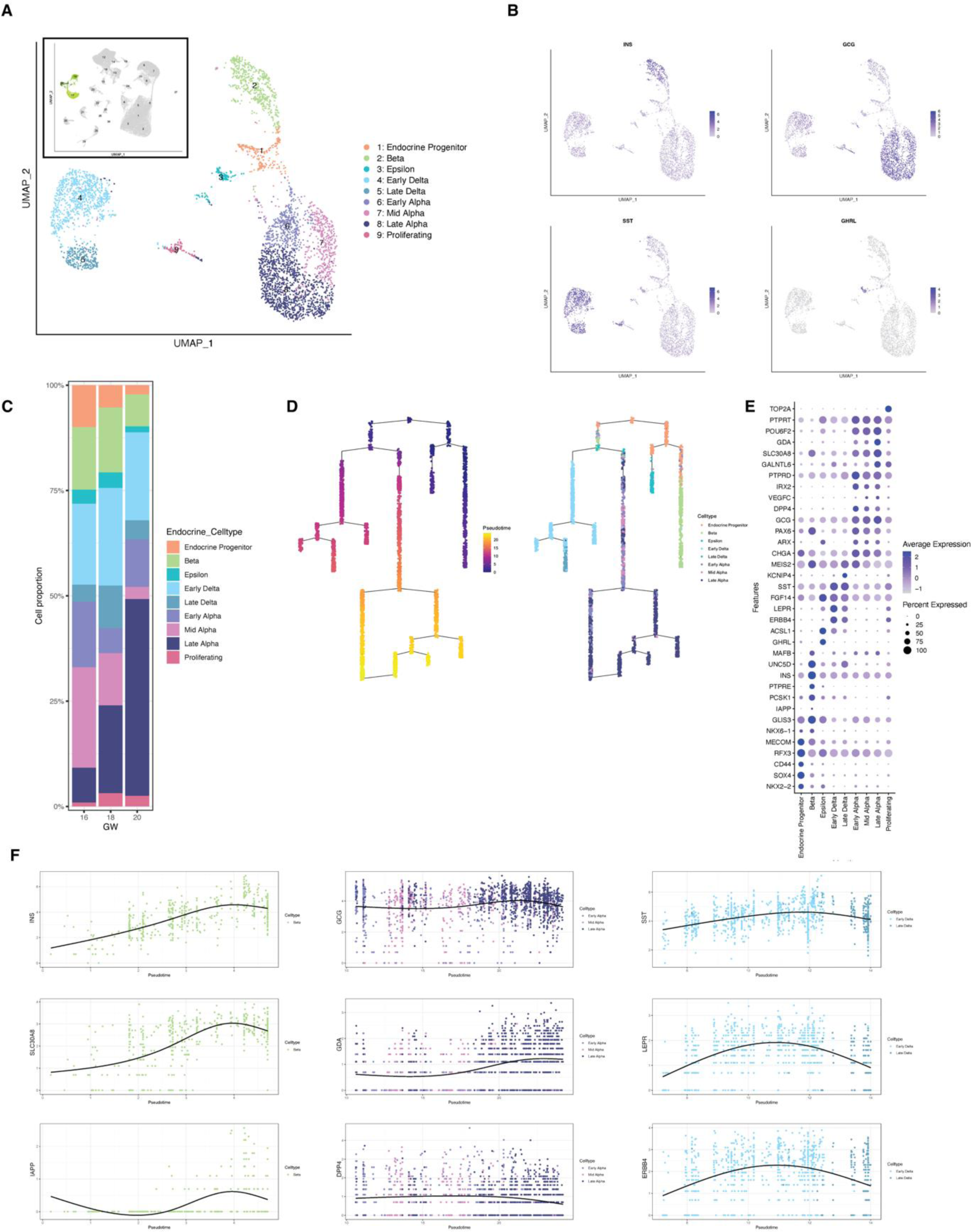
**A**,UMAP visualization and pseudotime-based cell annotation of pancreatic endocrine cells from Fig.1B. **B**, UMAP visualization of main endocrine hormones among pancreatic endocrine cells. **C**, Pancreatic fetal endocrine cell composition (frequency) by developmental stage. Colors indicate cell clusters as shown in panel A. **D**, Graph illustrating predicted trajectory of endocrine cells by pseudotime analysis. Colors of dots indicate a cell’s point in pseudotime (left) and cell annotation (right). **E**, Dot plot showing gene expression of endocrine cell markers. **F**, Scatterplot showing expression pattern of selected genes among alpha, delta and beta cells over pseudotime.

**Figure S7.**
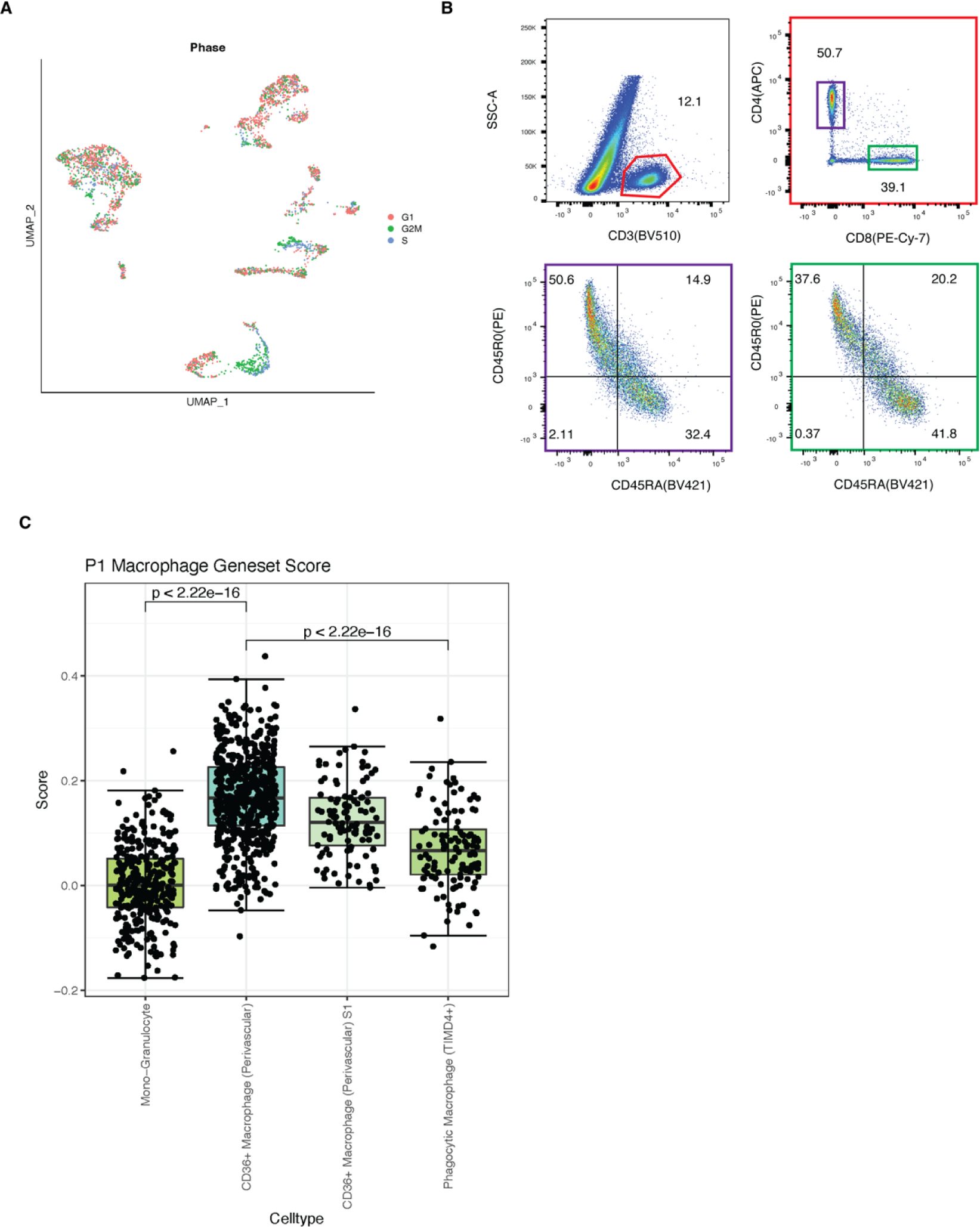
**A**,UMAP visualization of cell cycle analysis of fetal pancreatic hematopoietic cells. Colors indicate different phases of the cell cycle. **B**, Representative flow cytometry profile analysis of CD45RA+ and CD45RO+ among CD4+ and CD8+ T cells in fetal pancreas at 16 PCW. **C**, Boxplot showing comparison by canonical correlation analysis between the identified human fetal pancreatic macrophage subpopulations (this study) and P1 cardiac macrophages (data set from A.B Aurora et al., 2014)

**Figure S8.**
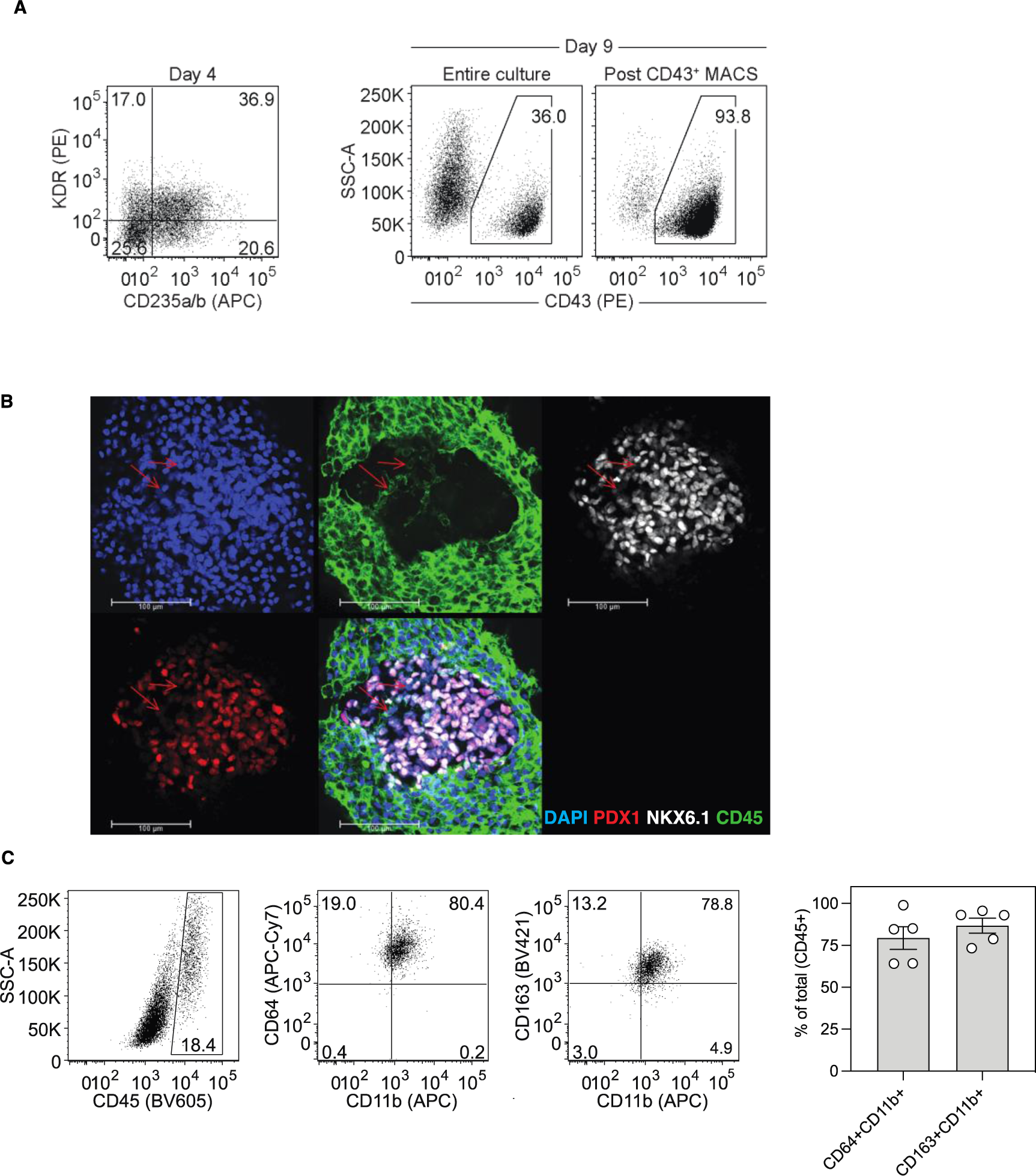
**A**,Flow cytometry profile of hESC-derived CD43+ embryonic macrophage progenitors. **B**, CLSM images of CD45+ (green) hESC-derived macrophages and (PDX1+NKX6.1+) endocrine organoids (eMAC-Endo) showing macrophages (in green) intermingling with the developing endocrine islets cells. **C**, Flow cytometry profile of hESC-derived eMAC phenotype upon 5 days in coculture with developing hESC-derived pancreatic progenitors.

**Figure S9.**
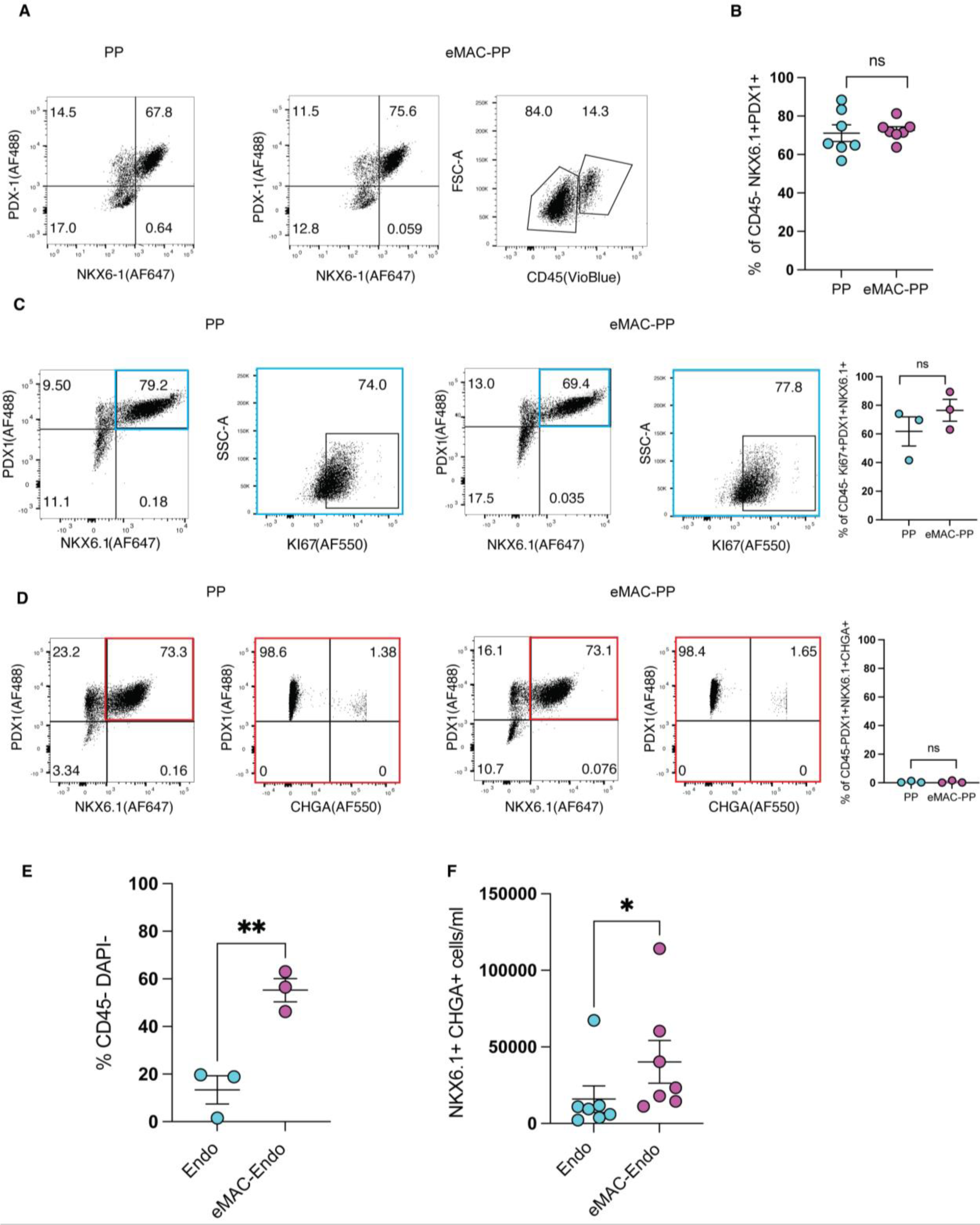
**A**,Flow cytometry profile of eMAC (CD45) and hESC-derived pancreatic progenitors (PDX1/NKX6-1) generated without (PP) and with (eMAC-PP) macrophages. **B**, Quantification of hESC-derived pancreatic progenitors generated without (PP) and with (eMAC-PP) macrophages (n=7 independent experiments, error bar represents SEM, unpaired t-test, ns: not significant) **C**, Flow cytometry profile and quantification of the proliferation rate of hESC derived pancreatic progenitors generated without (PP) and with (eMAC-PP) macrophages (n=3 independent experiments, error bar represents SEM, unpaired t-test, ns: not significant). **D**, Flow cytometry profile and quantification of CHGA within hESC derived PP generated without (PP) and with (eMAC-PP) macrophages (n=3 independent experiments, error bar represents SEM, unpaired t-test, ns: not significant). **E**, Quantification of viable endocrine islet cells generated without (Endo) and with (eMAC-Endo) macrophages using DAPI exclusion of live cells (n=3 independent experiments, error bar represents SEM, unpaired t-test, p= 0.0055). **F**, Quantification of endocrine progenitors generated without (Endo) and with (eMAC-Endo) macrophages (n=7 independent experiments, error bar represents SEM, paired t-test, p= 0.0159)

